# EVd3x: A Bioinformatics Platform for Evidence-aware Interpretation of EV Cargo

**DOI:** 10.64898/2026.05.06.723262

**Authors:** Karima Ait Ouares, Jonathan S. Weerakkody

## Abstract

Extracellular vesicle (EV) profiling can identify hundreds to thousands of RNAs and proteins, but converting those lists into a coherent biological interpretation remains slow because the relevant evidence is scattered across cargo repositories, publications, pathway databases, cell atlases, and interaction resources. We introduce EVd3x.com, a source-linked bioinformatics platform that brings these evidence layers into one continuous workflow for EV research. EVd3x harmonizes more than 22 million records and interactions from 28 public resources into 17 analysis tables, including 234,090 reported EV-cargo records linked to 3,350 publications, 2.6 million miRNA-target relationships, 404,758 pathway memberships, more than 501,000 disease associations, 1.6 million cell-context records, 25,779 ligand–receptor pairs, and 13.7 million protein-interaction records. Each query generates a fixed, exportable evidence packet that follows the same molecules from reported EV detection through disease, pathway, cellular, regulatory, and interaction analyses. Starting with the complex disease query “early-onset Alzheimer’s disease with behavioral disturbance,” EVd3x arrived at a focused, PSEN1-centered evidence packet and organized thousands of linked records into a source-traceable interpretation. The workflow recovered expected Alzheimer biology while keeping reported EV detection separate from broader contextual evidence, thereby exposing evidence gaps and context-dependent relationships. EVd3x provides EV researchers with a practical route from a cargo list, or from a disease, pathway, or cell-context question to a prioritized set of molecules, relationships, evidence gaps, and unresolved questions that researchers can use to formulate hypotheses and design subsequent experimental studies.

## 1. Introduction

Extracellular vesicles (EVs) can carry molecular information between cells [1] and are increasingly studied as biomarkers, mediators of intercellular communication, and therapeutic vehicles. Modern EV profiling can measure hundreds to thousands of RNAs and proteins in a single experiment. The result, however, is usually a molecular list rather than an explanation of what the molecules mean, which findings deserve attention, or what evidence must be verified next.

This interpretation problem is especially difficult in EV research because the biological meaning of a detected molecule depends on how the extracellular material was collected, enriched, characterized, and measured. EV preparations can contain heterogeneous vesicle populations together with lipoproteins, protein complexes, exomeres, supermeres, and other extracellular particles with overlapping physical properties [2–12]. A reported cargo therefore becomes useful only when it can be traced back to the sample, preparation method, detection technology, publication, and reporting quality that support it.

A second challenge begins after an EV-associated record is identified. Researchers must connect the candidate to disease biology, pathways, plausible source and recipient cells, miRNA targets, ligand-receptor relationships, and protein networks. Each resource captures a different part of this biological context, but these evidence layers are rarely reviewed together. The result is a fragmented workflow in which important connections may be missed, duplicated database records may appear stronger than they are, and successive analyses may unknowingly rely on different versions of the molecular list.

This fragmentation mirrors a broader challenge in EV research: evidence generated at different experimental stages must be connected without being treated as equivalent. The MISEV guidelines define minimum reporting and experimental requirements for EV studies, evolving from MISEV2014 [13] through MISEV2018 [14] to MISEV2023 [15]. EV-TRACK complements these guidelines by promoting transparent reporting [16] and linking experimental metadata to database records [17]. Additional standardization, EV-RNA, bioinformatics, urinary-EV, and reproducibility recommendations address sample handling, molecular localization, functional interpretation, and repeatability [18–23]. Together, these frameworks establish a practical evidence hierarchy in which preparation quality, cargo detection, molecular location, transfer, and function are considered connected but distinct stages of support. A useful computational platform should follow the same logic: preserve the original EV evidence, add biological context without losing provenance, identify where independent evidence converges or conflicts, and indicate the verification required to support, refine, or reject a proposed biological model.

We developed EVd3x.com to make this process continuous, scalable, and reproducible. The platform integrates more than 22 million records and interactions from 28 public resources into 17 harmonized analysis tables. These include 234,090 reported EV-cargo records from ExoCarta [24], Vesiclepedia [25], and SVAtlas [26], linked to 3,350 publication records and supplemented with EV-TRACK metadata where available [16,17]. Cross-resource identity mapping through Ensembl [27], UniProt [28], miRBase [29], and RNAcentral [30] allows genes, proteins, and miRNAs to move through the same analysis while remaining linked to their original source records.

Once molecular identities are harmonized, EVd3x adds the regulatory, pathway, and disease context needed to interpret those records biologically. This layer includes 2.6 million scored miRNA-target relationships derived from miRTarBase [31], DIANA-TarBase [32], and TargetScan [33]; 404,758 pathway memberships from Reactome [34], KEGG [35], Gene Ontology [36], and WikiPathways [37]; and more than 501,000 disease associations from DisGeNET [38].

To place these relationships within relevant tissues and cell types, EVd3x incorporates expression and localization data from the Human Protein Atlas [39], RNALocate [40], miRNATissueAtlas2 [41], and miRmine [42], together with 1.6 million harmonized cell-context records from the Human Protein Atlas [39], CellMarker [43], and PanglaoDB [44]. Building on this cellular context, the platform integrates 25,779 curated ligand-receptor pairs from CellPhoneDB [45,46], OmniPath Intercell [47], CellTalkDB [48], Cellinker [49], and Reactome [34], while 13.7 million protein-interaction records from STRING [50] provide broader network context. Together, these layers connect reported EV cargo to regulatory, disease, pathway, cellular, communication, and protein-network evidence while preserving each source as a distinct and traceable evidence layer. These counts represent a versioned snapshot rather than a complete catalogue of EV cargo. As new EV studies, cargo records, and EV-TRACK metadata become available, EVd3x is designed to incorporate the expanding knowledge base while preserving source provenance and database versioning. (Methods: Canonical table construction and source harmonization; Supplementary Note 1; Figures S1–S2; Table S1).

EVd3x differs from a collection of independent database searches in one central respect: every query is converted into a fixed, reviewable evidence packet. Direct inputs, inferred molecules, source-linked records, filters, thresholds, and processing limits are preserved as the analysis moves from reported EV evidence to disease, pathway, cell-context, communication, miRNA-target, and protein-interaction layers. By carrying the same molecular state through each step, EVd3x prevents unnoticed changes in the analyzed list, makes the resulting biological interpretation reproducible, and allows every displayed result to be exported and traced to the evidence that produced it.

EVd3x is intended to support both cargo-first and biology-first analysis. Users may begin with an experimental cargo list or individual molecules, or with a disease, pathway, cell type, communication setting, or natural-language question and use the returned records to construct a manageable molecular packet. As a worked example, we entered the complex query “early-onset Alzheimer’s disease with behavioral disturbance.” The query led to the PSEN1-centered evidence packet recorded in Table S2, providing a biologically recognizable positive-control setting because PSEN1, APP processing, γ-secretase and Notch biology, APOE, BACE1, neuronal and glial states, and Alzheimer-associated EV studies are extensively described [51–67]. The case study therefore asked whether an integrated EV platform could carry one evidence packet through reported EV, disease, pathway, cellular, regulatory, and interaction layers without losing provenance, and whether the resulting record could identify candidates, conflicts, and verification requirements for subsequent testing. The source-attributed EV evidence for this packet is summarized in Figure S3.

Figures 1–6 follow the evidence-review sequence used in the case study. The workflow begins with the early-onset Alzheimer query and its PSEN1-centered evidence packet, then examines reported EV-cargo support, disease and pathway context, candidate cellular settings, and regulatory and interaction relationships. Figure S4 maps each result to its source records, Table 1 defines what each evidence layer can and cannot support, and Tables S1–S10 provide the row-level exports, parameters, and sensitivity analyses. Together, these materials show how EVd3x turns a broad starting question or experimental cargo list into a prioritized, traceable set of candidate molecules, relationships, evidence gaps, and questions requiring independent verification.

**Table 1.** Evidence hierarchy for interpreting EVd3x outputs and identifying requirements for experimental verification. Note: Each layer supports a different level of interpretation. Evidence-aware use begins with reported EV detection and study provenance; Broader annotations provide context that researchers may use when deciding what to investigate; they do not determine an experimental plan or replace independent validation. [14–17].

| **EVd3x evidence layer** | **What it can support** | **What it cannot support** | **Examples of independent verification** |
| --- | --- | --- | --- |
| **EV cargo evidence row** | Reported detection of a molecule in an EV or EV-enriched resource/publication; source, sample, method, PubMed, and EV-TRACK-linked context when available | True vesicular loading, EV subtype, preparation purity, abundance, disease specificity, uptake, or function | Orthogonal EV isolation; EV-marker characterization; particle/protein ratio; detergent/protease protection where appropriate; qPCR/ddPCR, Western blot, or MS validation |
| **EV-TRACK/reporting metadata** | Availability of study-level reporting and protocol information | A guarantee of experimental quality or functional validation | Review the original publication, isolation method, markers, controls, and reporting completeness |
| **Disease association** | Disease-context annotation for genes, proteins, or miRNAs | Biomarker validity, diagnostic performance, or causality | Independent cohort testing; disease-control comparisons; ROC and validation studies |
| **Pathway enrichment** | Over-representation of pathway annotations among query or expanded molecules | Pathway activity in EV-producing or recipient cells | Perturbation assays; protein/RNA validation; pathway reporter or functional assay |
| **Cell-context ranking** | Candidate tissue or cell-context based on marker and expression resources | EV cell of origin, selective cargo packaging, or uptake | Cell-specific EV enrichment or immunocapture; single-cell or spatial validation; cell-specific perturbation |
| **Ligand-receptor evidence** | A candidate communication pair reported by curated ligand-receptor resources | EV-surface display, EV-mediated transfer, receptor activation, or in vivo direction | Verify molecular roles and membrane location; validate surface exposure; use receptor blocking, transfer assays, or co-culture perturbation |
| **miRNA-target bridge** | A candidate regulatory relationship from experimental or predicted target resources | EV delivery, target repression, or functional regulation in the studied system | Luciferase assay; miRNA perturbation; qPCR and protein validation in recipient cells |
| **STRING/PPI edge** | Protein-interaction or association context | An EV-specific interaction or pathway activation | Co-IP or proximity assay; perturbation; pathway readout |

| EVd3x evidence layer | What it can support | What it cannot support | Examples of independent verification |
| --- | --- | --- | --- |
| **Assistant summary** | Navigation and explanation of retrieved information, table-backed evidence | Independent biological evidence, expert adjudication, or a new discovery | Inspect exported rows and the original source publications |

## 2. Results

The Results follow the order in which an EV researcher would review a candidate set: establish a stable molecular state, inspect reported EV support, place the state in disease and pathway context, evaluate candidate cellular settings, and then review regulatory and interaction relationships. At each step, EVd3x preserves the source records and the claims that the evidence can, and cannot, support.

### 2.1 A fixed Alzheimer case study demonstrates the EVd3x evidence workflow

The exact Alzheimer query was entered without a user-supplied molecule list. EVd3x recognized it as a disease-oriented request and mapped it to familial Alzheimer disease type 3. The query led to the PSEN1-centered case-study packet reported in Table S2. For manuscript reproducibility, this packet was held constant across all downstream analyses. Its visible starting molecules were PSEN1 mRNA, its corresponding protein representation, and three miRNAs: hsa-miR-4496, hsa-miR-107, and hsa-miR-4677-3p, from the case-study candidate set (Figure 1A–B; Methods: Query workflow and reviewable evidence packet; Supplementary Notes 2–3; Figure S5; Table S2). These five molecules served as the visible starting points for review across the platform.

**Figure 1.**
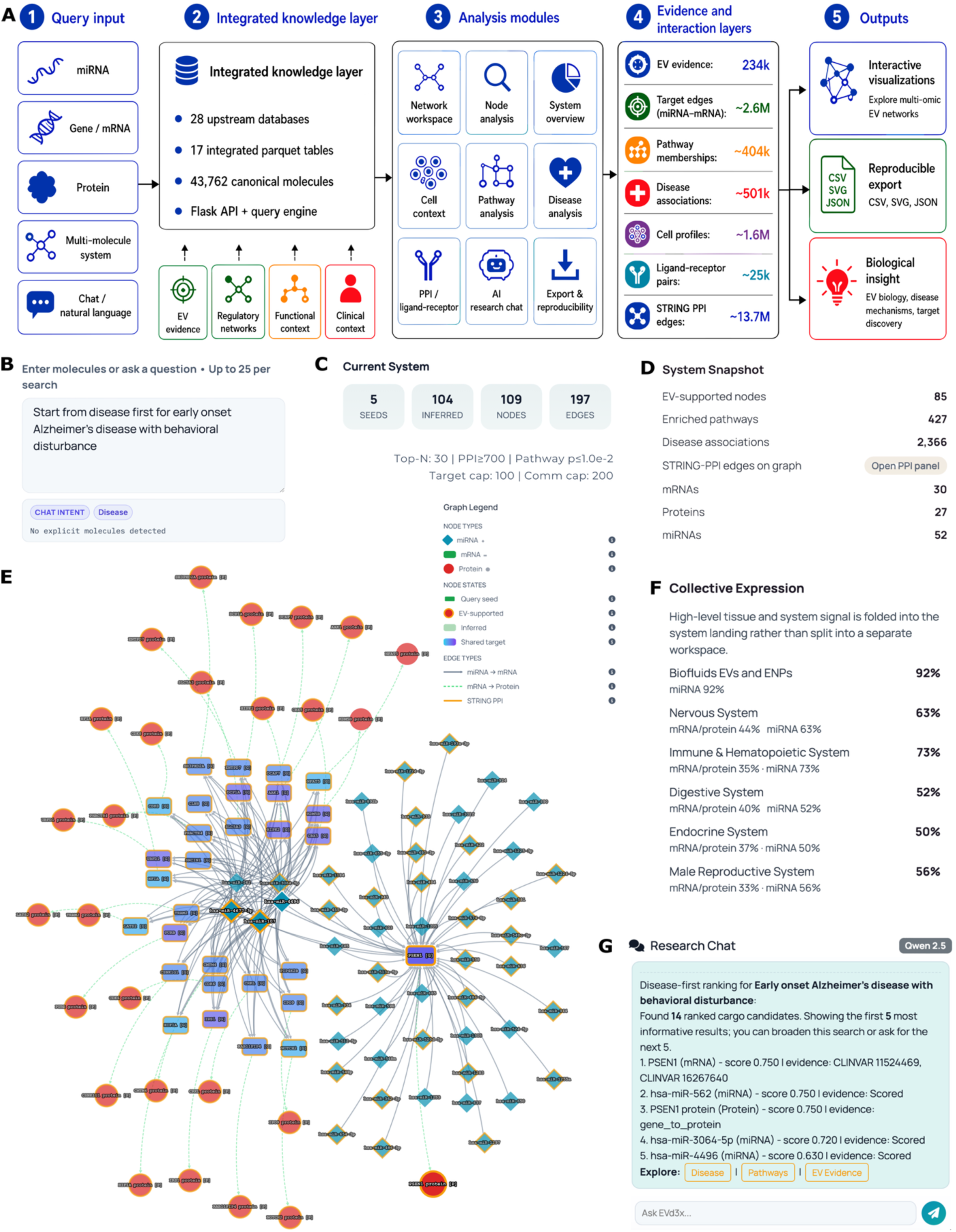
A disease-first question is resolved into a fixed, reviewable evidence packet. **A)** Schematic representation of the EVd3x workflow, showing how individual miRNAs, genes or mRNAs, proteins, molecular collections, and natural-language questions enter a shared evidence workflow. **B)** The Alzheimer query is entered without an explicit molecule identifier; deterministic routing and table retrieval lead to the PSEN1-centered case-study packet before optional language-model synthesis. **C)** The fixed workspace contains five visible starting nodes, 104 inferred molecules, 109 total nodes, 197 displayed relationships, and the active processing limits. **D)** The same evidence packet summarizes nodes with reported EV support, pathway and disease coverage, and molecule classes. **E)** The graph displays molecule and relationship classes for evidence navigation. **F)** Expression and system-level summaries identify candidate cellular settings for review. **G)** The assistant summarizes the retrieved records without becoming an additional evidence source. The workflow is described in Methods: Query workflow and reviewable evidence packet, and Natural-language navigation and grounding. Reproducibility parameters and source mappings are provided in Supplementary Notes 1–3, Figure S5, and Tables S1–S2.

The first-pass display contained 14 ranked candidates spanning direct PSEN1 anchors, PSEN1-targeting miRNA bridges, recommended miRNAs, and Alzheimer-relevant network neighbors. Three additional miRNAs: hsa-miR-562, hsa-miR-3064-5p, and hsa-miR-1184, were retained because the integrated miRNA–target table linked them to PSEN1. The fixed gene universe contained PSEN1 plus 25 target-derived or network-context genes, including APP, BACE1, APOE, MAPT, PSEN2, NOTCH-family genes, CTNNB1, and several kinase-associated candidates (Supplementary Notes 2 and 7; Tables S2 and S7). Each molecule remained labeled by its inclusion route so that direct anchors, regulatory bridges, recommendations, and network-context candidates could be distinguished.

The fixed Alzheimer packet contained five visible starting molecules, 104 inferred molecules, 109 total molecular nodes, and 197 displayed relationships (Figure 1C; Table 2; Methods: Query workflow and reviewable evidence packet; Table S2). These counts define the reproducible state carried through subsequent analyses; they do not measure evidence strength, independence, or biological importance. Because every module read from the same state, each result could be traced to the recorded molecule set, source rows, filters, thresholds, and processing settings.

Within this packet, 85 nodes had reported EV-associated support, and the active state intersected 427 pathway summaries and 2,366 disease-association summaries (Figure 1D–F; Methods: Canonical table construction and source harmonization; Supplementary Notes 1–2; Figures S1 and S4; Tables S1–S2). These counts describe annotation breadth. Well-studied or highly connected molecules generate more records, so graph position, connectivity, and record count were used for navigation rather than as measures of biological importance.

The fine-tuned language model was invoked only after deterministic intent routing and table-based retrieval had produced the fixed packet and its source-linked records (Figure 1G; Methods: Natural-language navigation and grounding; Supplementary Notes 9–11; Figure S6; Table S8). It summarized the retrieved evidence and highlighted candidates for review, but it did not select the starting molecules, calculate scores, create biological associations, or add network edges. Figure 1 therefore defines the central analytical unit of the case study: a fixed, inspectable, exportable, and reproducible packet that can be followed across evidence layers without losing provenance.

**Table 2.** Scale of the integrated resource and the fixed Alzheimer case-study evidence packet. Counts describe database records, retrieved evidence rows, or displayed summaries rather than evidence or validation strength. Source inventories and the complete case-study parameters are provided in Tables S1–S2.

| Feature | Footprint |
| --- | --- |
| **Source integration** | 28 public resources resolved into 17 canonical Apache Parquet analysis tables. |
| **Evidence layers** | Identity, reported EV cargo, publication and EV-TRACK metadata, miRNA–target, pathway, disease, expression, localization, cell-context, ligand–receptor, and protein-interaction records retained as distinct evidence layers. |
| **Case study entry** | Manuscript-specific natural-language preset: “early-onset Alzheimer’s disease with behavioral disturbance.” |
| **Case-study molecular state** | 5 visible starting nodes comprising PSEN1 mRNA, its corresponding protein representation, and 3 first-pass miRNA candidates; 3 additional PSEN1-targeting miRNAs retained for bridge analysis; and 25 target-derived or protein-network context genes. |
| **Row-level support** | 647 reported EV-evidence rows, 4,053 disease-association rows, 2,204 pathway-membership rows, 3,555 cell-context, communication, and ligand–receptor rows, and 4,032 miRNA-target and protein-interaction bridge rows. |
| **Interactive processing guard** | The live interface accepts up to 25 input molecules per graph request. Requests exceeding this limit are blocked under the default release and must be divided into separate searches or processed through a separately configured workflow. This is a computational limit, not a biological threshold. |

### 2.2 Source-linked records define the reported EV-cargo evidence boundary

To establish the EV-specific evidence boundary, we asked which source-linked records reported PSEN1 in EV or EV-enriched material. Within the fixed case-study state, PSEN1 protein was selected as the focal molecule and anchored to the reviewed UniProt accession P49768. Its molecular identity and reported EV-associated records were examined separately from target-derived candidates and protein-network neighbors (Figure 2A–B; Methods: EV evidence retrieval and reported-cargo provenance; Supplementary Note 4; Table S3).

**Figure 2.**
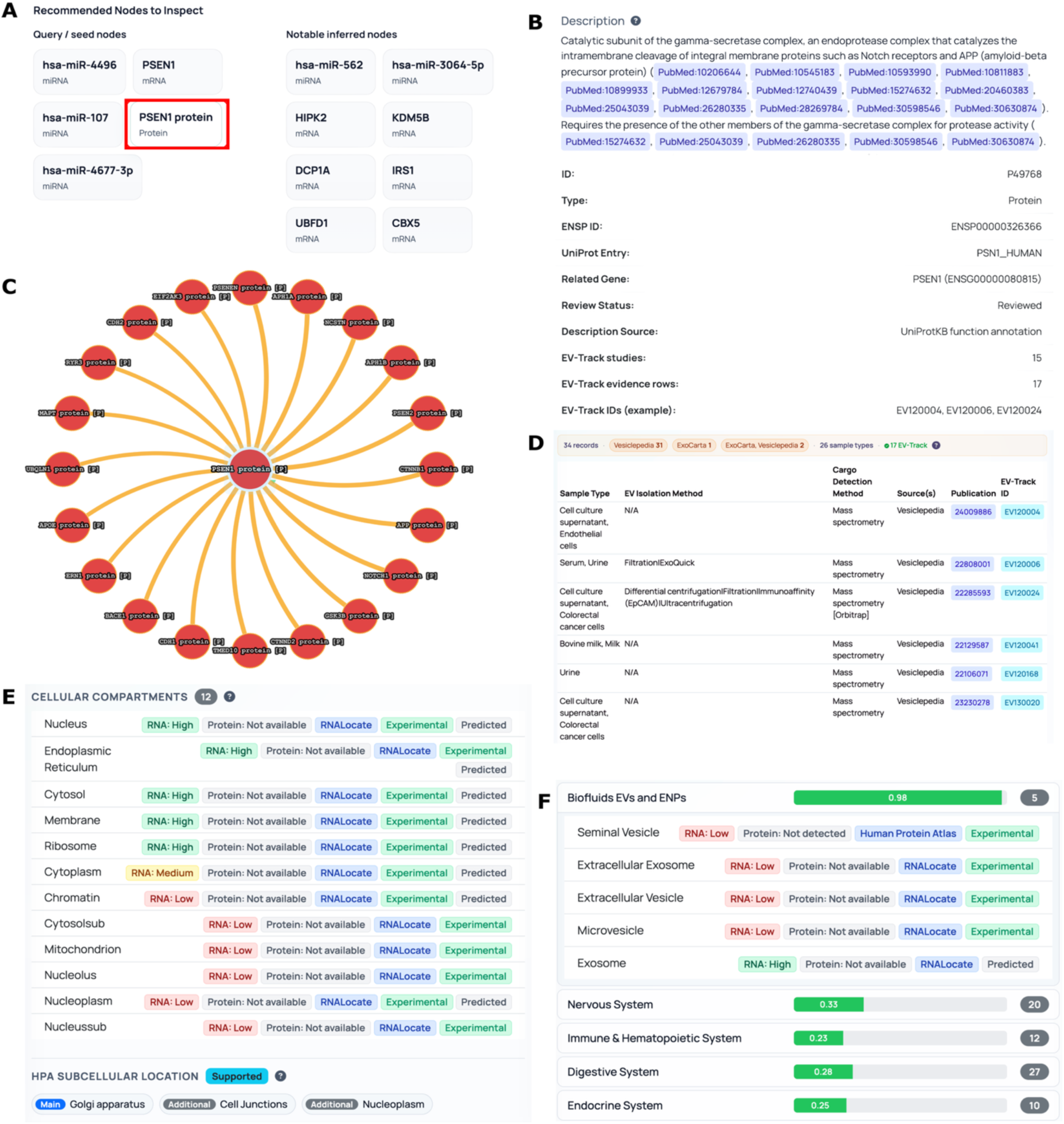
PSEN1 review separates molecular identity and network context from reported EV provenance. **A)** Visible starting molecules are separated from inferred candidates, and PSEN1 protein is selected as the focal molecule for review. **B)** The detail view anchors PSEN1 to reviewed molecular identifiers and γ-secretase biology. **C)** The high-confidence protein neighborhood provides APP-processing, Notch, and neurodegenerative context but is not treated as reported EV evidence. **D)** Thirty-four displayed reported EV-cargo records for PSEN1, including 17 EV-TRACK-linked records, retain source, sample, preparation, detection, publication, and reporting fields. **E)** Localization records provide cellular-compartment context. **F)** Biofluid and system annotations retain extracellular-vesicle labels for review and export. EV-evidence retrieval and PPI filtering are described in Methods: EV evidence retrieval and reported-cargo provenance, and miRNA-target and PPI bridge prioritization. Source-level and case-study packet audits are provided in Supplementary Note 4, Figures S2–S3 and S7C, and Tables S3 and S7. The figure supports review of reported detection and experimental context; selective loading, intravesicular localization, abundance, disease specificity, and function require independent verification.

The high-confidence protein neighborhood placed PSEN1 within APP-processing, γ-secretase, Notch, APOE/MAPT, and kinase-associated contexts (Figure 2C; Methods: miRNA-target and PPI bridge prioritization; Supplementary Note 8; Figure S7C; Table S7). This network provided the expected Alzheimer-related context while leaving reported EV evidence restricted to the source-attributed cargo records.

Thirty-four reported EV-cargo rows for PSEN1 were retrieved from the integrated ExoCarta, Vesiclepedia, and SVAtlas table; 17 were linked to EV-TRACK metadata (Figure 2D; Methods: EV evidence retrieval and reported-cargo provenance; Supplementary Note 4; Figures S2–S3; Table S3). The records spanned multiple sample types, preparation procedures, detection technologies, and publications. Some came from Alzheimer-related or neural biofluid studies [63–67], whereas others came from broader proteomic or cell-line investigations. Retained source, publication, and EV-TRACK fields allowed the experimental context of each record to be reviewed directly.

These rows establish that PSEN1 has been reported in EV or EV-enriched material across several experimental settings. Because the underlying studies differ in sample type, preparation, and assay, the records do not establish selective packaging, molecular topology, abundance, disease specificity, cellular origin, transfer, or function. Records must also be collapsed by PubMed or EV-TRACK study identifier before they are counted as independent studies.

Localization and biofluid annotations further identified the cellular compartments, sample settings, and publications associated with PSEN1 (Figure 2E–F; Methods: Canonical table construction and source harmonization; Supplementary Notes 1 and 4; Tables S1 and S3). These annotations help define the settings most relevant for follow-up, including independent confirmation of PSEN1 in an appropriately characterized preparation and, where relevant, determination of whether the signal is intravesicular or surface-associated. Functional interpretation would require separate evidence of transfer, target engagement, or biological response.

### 2.3 Disease associations recover the expected Alzheimer context

After reviewing the reported EV-cargo records, we examined how the fixed PSEN1-centered packet mapped across disease-association resources. Nervous-system disease categories dominated the returned disease space, and familial Alzheimer disease type 3 appeared at the top of the fixed case-study ranking (Figure 3A–C; Methods: Disease and pathway annotation; Supplementary Note 5; Figures S7D and S8B; Table S4). This provided the expected disease context for a PSEN1-centered positive-control analysis.

**Figure 3.**
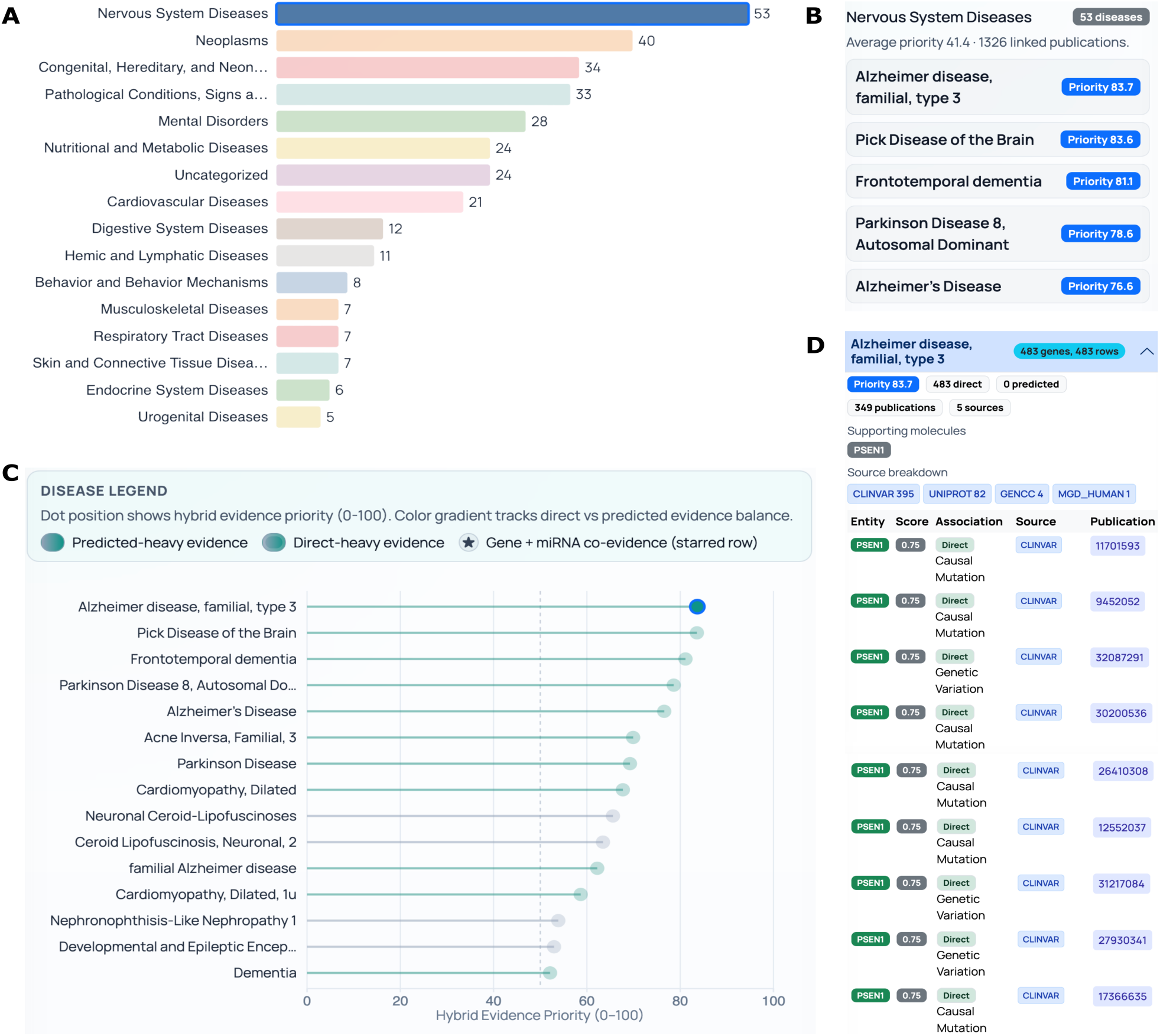
Disease associations place the fixed PSEN1-centered packet within the expected Alzheimer context. **A)** Disease-category counts summarize the returned association space, which is dominated by nervous-system disease labels. **B)** Disease cards display familial Alzheimer disease type 3 and related neurodegenerative conditions in the fixed case-study order. **C)** The ranking shows the review-priority score together with the balance of direct and predicted evidence. **D)** The selected familial Alzheimer disease type 3 record contains 483 direct association rows linked to 349 publications and five source categories, with PSEN1 as the supporting molecule. Disease aggregation and ranking are described in Methods: Disease and pathway annotation. Source composition and the authoritative row-level records are provided in Supplementary Note 5, Figures S7D and S8B, and Table S4. The ranking organizes evidence review according to curated evidence burden, source breadth, directness, and literature density; it should not be interpreted as disease probability, biomarker performance, or causal evidence.

The selected disease record contained 483 direct association rows linked to 349 publications and five source categories, with PSEN1 as the supporting molecule (Figure 3D; Table S4). These source-attributed records connected the evidence packet to the established familial Alzheimer literature and provided a direct route from the summarized result to the underlying associations and publications.

The scale of this record set reflected the depth of existing curation and literature coverage. Repeated publications and overlapping database sources can contribute multiple rows to the same underlying relationship, while extensively studied genes such as PSEN1 are represented more densely than less characterized candidates. The priority score therefore organized review order by combining evidence burden, supporting-entity burden, source and publication breadth, direct-evidence fraction, and source score; it was not interpreted as disease probability, biomarker performance, or causal strength.

The disease layer therefore showed that the fixed evidence packet was situated within the intended Alzheimer-associated literature and identified the records contributing to that context. It complemented the reported EV-cargo review in Section 2.2 but did not add evidence that PSEN1 was disease-specific or differentially represented in Alzheimer-associated EVs.

### 2.4 Pathway convergence organizes annotation breadth into candidate biological themes

After defining reported EV evidence and disease context, we asked whether the molecules in the fixed packet converged on recognizable biological pathways. Hypergeometric enrichment with Benjamini–Hochberg correction identified 153 manuscript-visible terms across 16 categories from Gene Ontology, KEGG, Reactome, and WikiPathways (Figure 4A; Methods: Disease and pathway annotation; Supplementary Note 5; Figures S7A and S9; Table S5). The source-specific contributions and large uncategorized component show that this landscape is shaped by database coverage, pathway definitions, and the selected filtering thresholds.

**Figure 4.**
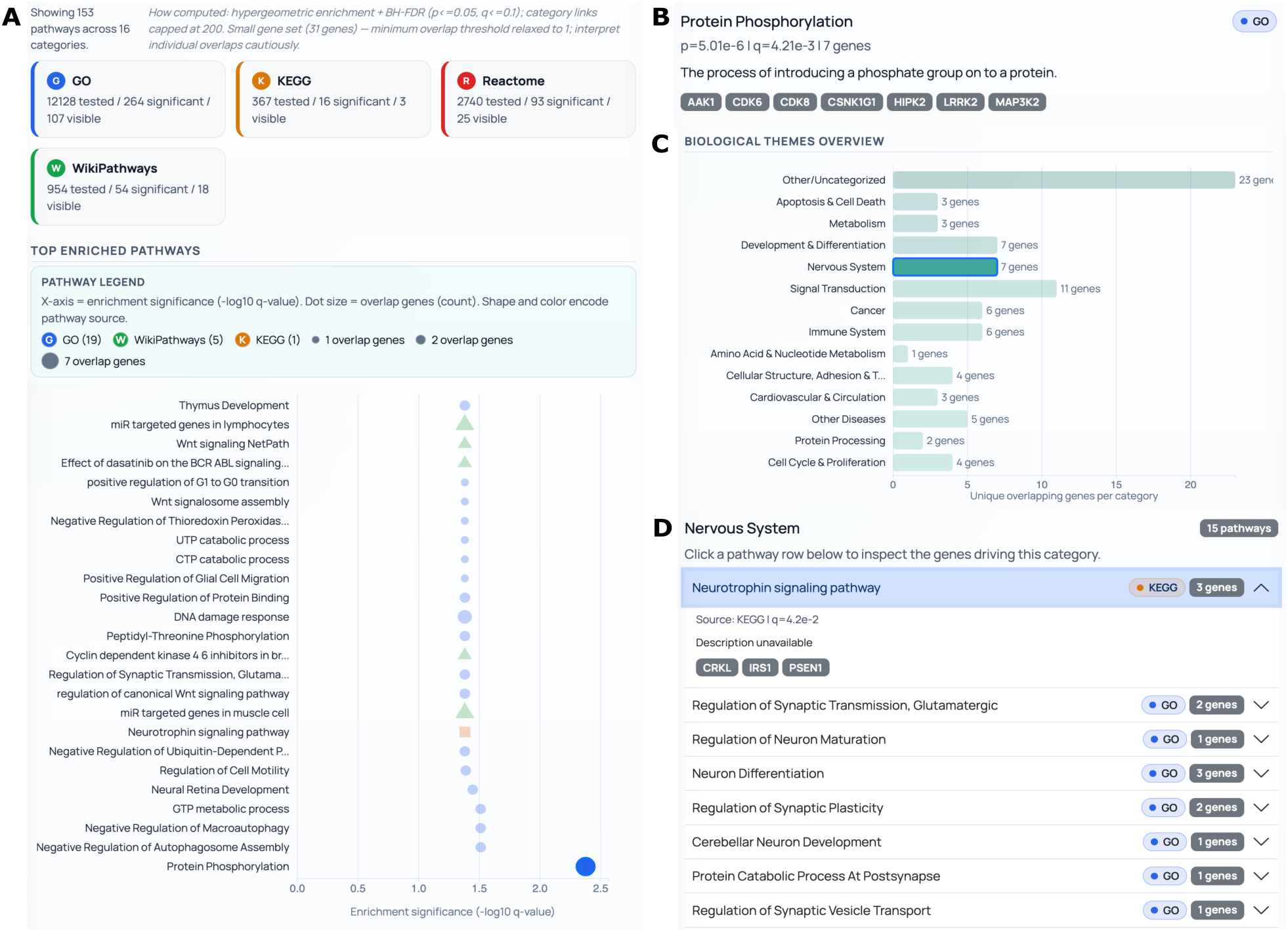
Pathway annotations organize the fixed evidence packet into candidate biological themes. **A)** Hypergeometric over-representation analysis with Benjamini–Hochberg correction identifies 153 manuscript-visible terms across Gene Ontology, KEGG, Reactome, and WikiPathways. **B)** Protein phosphorylation illustrates a kinase-associated theme supported by seven overlapping genes. **C)** Individual pathway terms are grouped into broader biological categories while retaining the large uncategorized component. **D)** Nervous-system terms highlight neurotrophin, synaptic, neuronal-maturation, and vesicle-transport contexts. Pathway enrichment and category construction are described in Methods: Disease and pathway annotation. Source contributions, threshold behavior, and authoritative row-level memberships are provided in Supplementary Note 5, Figures S7A and S9, and Table S5. These results organize biological context and review priorities; they do not demonstrate pathway activity in EV-producing cells, EV preparations, or recipient cells.

Rather than interpreting the 153 terms as independent findings, EVd3x organized them into a smaller number of recurring biological themes. One theme linked presenilin and Notch biology with APP and BACE1 processing; a second contained kinase- and phosphorylation-related genes, including AAK1, CDK6, CDK8, CSNK1G1, HIPK2, LRRK2, and MAP3K2; and a third connected neurotrophin, synaptic, neuronal-maturation, and vesicle-transport annotations (Figure 4B–D; Table S5).

This organization reduced a long enrichment table to a manageable set of candidate biological themes. APP-processing proteins, kinase-associated relationships, and neurotrophin or synaptic pathways emerged as focused areas for review. These themes prioritize where to look next; they do not demonstrate that a pathway is active.

Over-representation indicates that genes in the evidence packet occur more frequently than expected within particular annotation sets. It does not establish pathway activity in the EV-producing cell, representation of the relevant pathway components in the EV preparation, or a response in a recipient cell. These pathway themes therefore provide biological context and guide prioritization, but they require independent confirmation in the relevant EV and cellular setting.

Disease associations and pathway enrichment therefore served complementary roles: one placed the packet within the expected Alzheimer literature, and the other organized its molecular content into coherent biological themes. Neither changed the strength of the reported EV-cargo evidence established in Section 2.2.

### 2.5 Cell-context analysis identifies candidate settings for further evaluation

We next examined which cellular contexts were most strongly represented within the fixed evidence packet. Oligodendrocyte-lineage and other nervous-system cell labels ranked highly, while immune and other potentially confounding contexts remained visible (Figure 5A; Methods: Cell-context and communication prioritization; Supplementary Note 6; Figure S8C; Table S6). Retaining these alternatives is important because public expression resources, tissue datasets, and plasma-associated EV records may reflect mixed or overlapping cellular sources.

**Figure 5.**
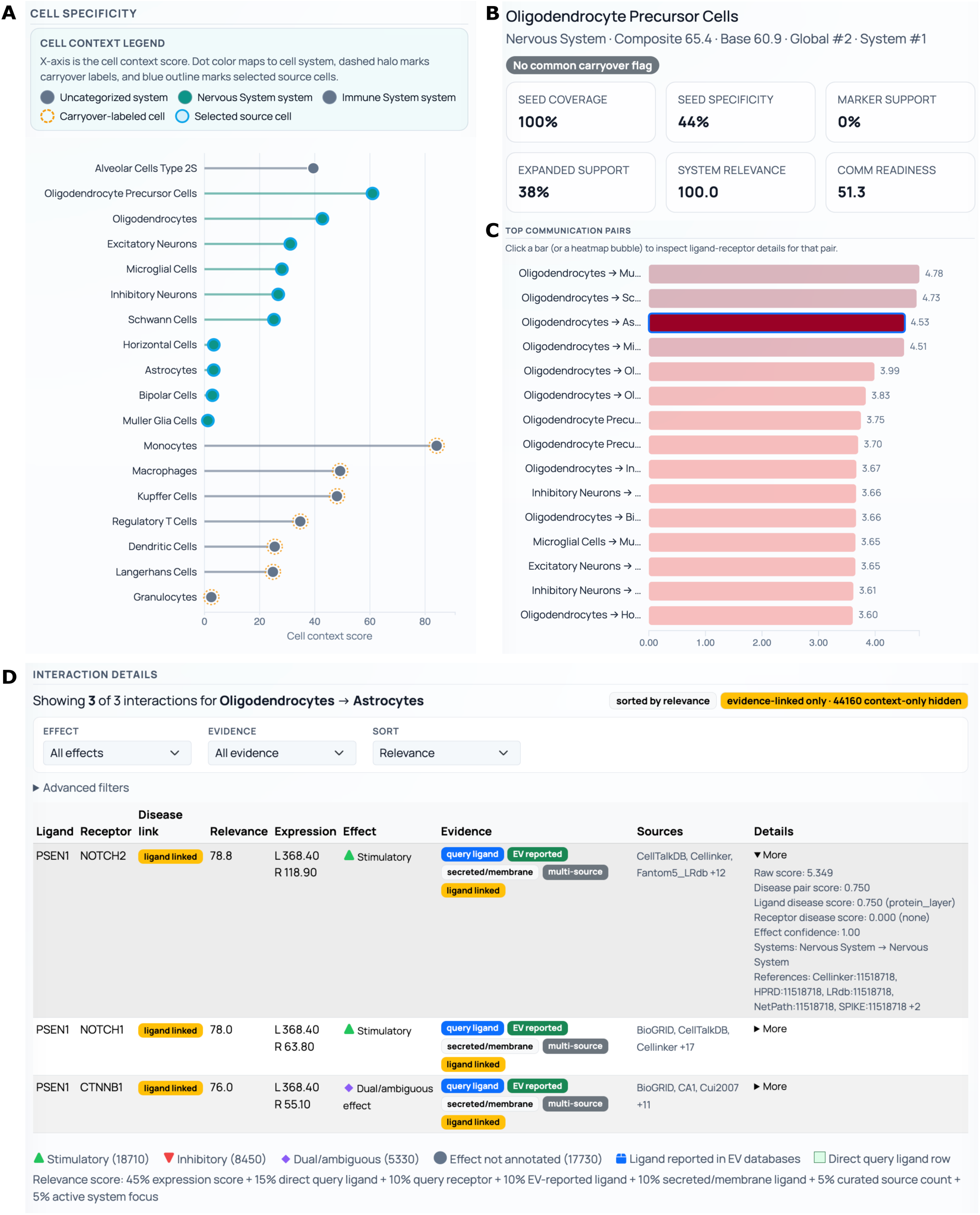
Cell-context and communication rankings are interpreted through their component evidence. **A)** Cell-context ranking retains nervous-system, immune, and other candidate settings for review. **B)** The oligodendrocyte precursor cell card shows that its high rank is driven by complete seed coverage and nervous-system relevance, despite moderate seed specificity, limited expanded-gene support, and no marker support. **C)** The communication view examines an oligodendrocyte-to-astrocyte setting. **D)** PSEN1–NOTCH2, PSEN1–NOTCH1, and PSEN1–CTNNB1 records remain after context-only relationships are hidden. Cell-context and communication scoring are described in Methods: Cell-context and communication prioritization. Component evidence and source-level audits are provided in Supplementary Note 6, Figures S7B and S8C–D, and Table S6. These outputs identify candidate cellular and communication settings for further review; they do not establish EV cell of origin, uptake, directionality, surface presentation, or receptor activation.

The oligodendrocyte precursor cell result illustrates why the score components are more informative than the overall rank alone. The selected card showed 100% seed coverage and complete nervous-system relevance, but only 44% seed specificity, 38% expanded-gene support, and no marker support, with a communication-readiness score of 51.3 (Figure 5B; Table S6). Its high rank was therefore driven primarily by query coverage and system-level weighting rather than by strong cell-marker evidence.

Accordingly, the result is best interpreted as a candidate cellular context rather than evidence that the queried cargo originated from oligodendrocytes. The ranked outputs can instead be used to compare plausible settings, including oligodendrocyte-lineage, astrocyte, neuronal, and immune models, and to identify the evidence needed to distinguish among them, such as cell-specific EV enrichment, immunocapture, donor-cell perturbation, single-cell or spatial data, or direct comparison of candidate source cells.

The communication view then examined an oligodendrocyte-to-astrocyte context and retained PSEN1–NOTCH2, PSEN1–NOTCH1, and PSEN1–CTNNB1 records after context-only relationships were hidden (Figure 5C–D; Methods: Cell-context and communication prioritization; Supplementary Note 6; Figures S7B and S8D; Table S6). These records show that the selected molecules and curated interaction labels coexist within the chosen cell-pair context. Their interpretation, however, depends on the assigned molecular roles, directionality, membrane localization, and relevance to EV-associated communication.

Together, the cell-context and communication analyses narrowed the candidate cellular settings for review while exposing weak marker support and uncertain molecular roles that must be resolved before advancing cell-of-origin or communication claims.

### 2.6 Integrated relationship evidence identifies candidates and rejection checks

The final analysis combined ligand–receptor, miRNA–target, and protein-interaction records to identify molecular relationships that warranted further review. The ligand–receptor workspace contained 323 candidate pairs, and PSEN1–CTNNB1 was selected as an example of a direct-overlap relationship within the active packet (Figure 6A–B; Methods: Cell-context and communication prioritization; Supplementary Note 8; Figure S7B; Table S6).

**Figure 6.**
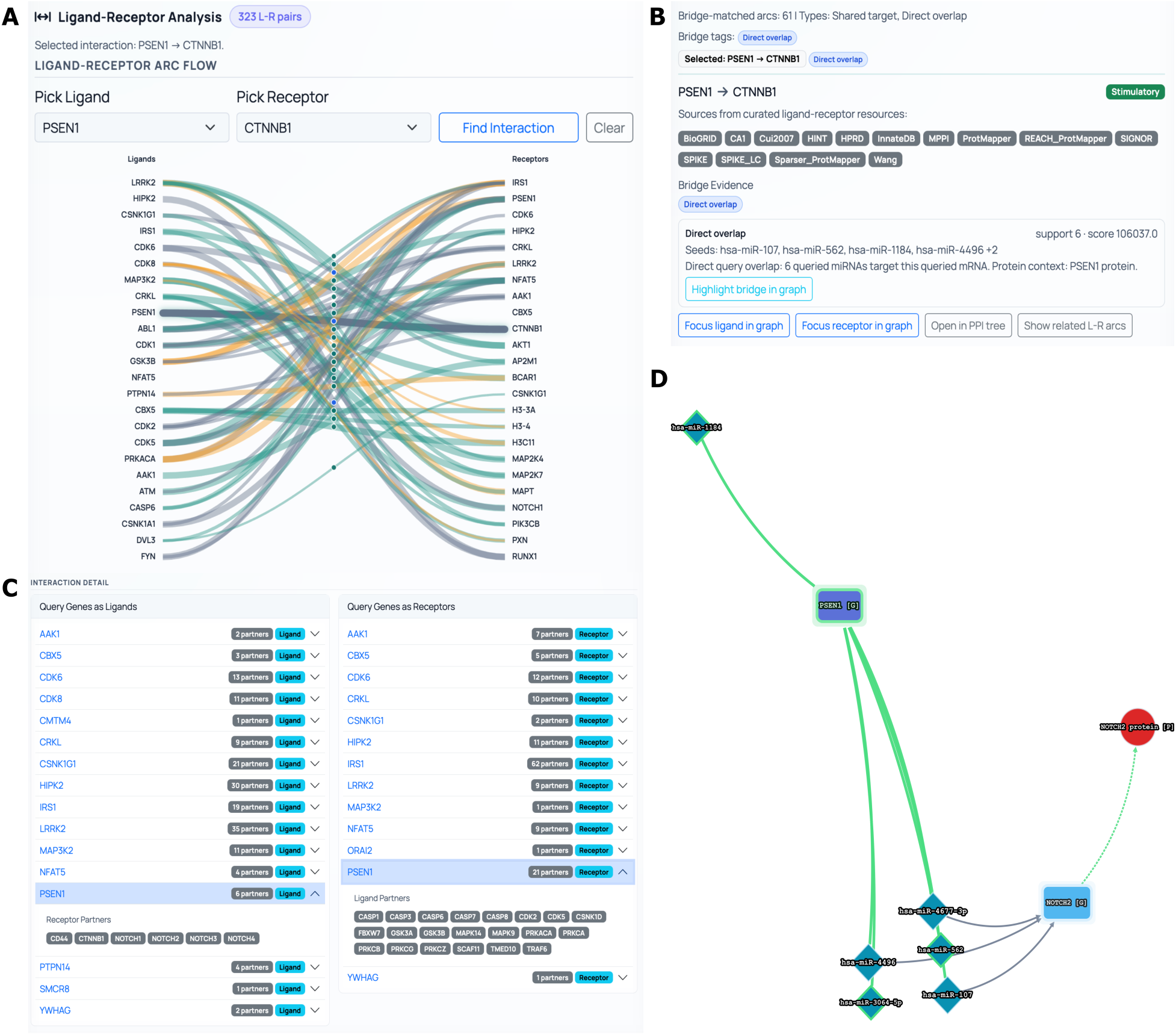
Integrated relationship layers identify candidates and rejection checks. **A)** The ligand–receptor workspace displays 323 candidate pairs and highlights PSEN1–CTNNB1 within the active evidence packet. **B)** The selected relationship retains curated source labels and active-state overlap while exposing a molecular-role assignment that requires biological review. **C)** Role-specific tables preserve source identifiers, scores, evidence classes, and partner information for inspection and export. **D)** Six miRNAs retained in the case-study packet have target support for PSEN1, while miR-107 also connects the packet to BACE1 and APP-processing biology. Ligand– receptor, miRNA-target, and protein-interaction procedures are described in Methods: Cell-context and communication prioritization, and miRNA-target and PPI bridge prioritization. Source composition and threshold audits are provided in Supplementary Notes 7–8, Figures S7B–C, and Tables S6–S7. These relationship layers prioritize records for further review and expose reasons to clarify or reject them; they do not establish EV localization, transfer, target repression, receptor activation, or pathway function.

The PSEN1–CTNNB1 example also exposed an important point of uncertainty. PSEN1 is an integral membrane catalytic component of γ-secretase rather than a conventional soluble ligand, while PSEN1–NOTCH1/2 records may reflect pathway or interaction annotations rather than demonstrated ligand–receptor communication. The originating source, assigned molecular roles, membrane localization, directionality, and experimental context must therefore be reviewed before these records are advanced as communication candidates.

The miRNA–target layer identified a different form of convergence. Six miRNAs retained in the case-study packet had target support for PSEN1, while miR-107 also connected the packet to BACE1 and APP-processing biology [56] (Figure 6B–D; Methods: miRNA-target and PPI bridge prioritization; Supplementary Note 7; Table S7). These relationships were prioritized for targeted review and measurement, while their EV localization, delivery, and functional regulation remained subject to independent verification.

Table 1 and Supplementary Table S9 place these relationships within a three-level verification hierarchy. The first level addresses candidate detection and molecular location in an appropriately characterized EV preparation; the second addresses cellular context and transfer; and the third addresses target engagement or mechanism. The framework identifies the evidence required to advance each relationship without implying that those experiments were performed or that EVd3x independently validated the proposed biology.

The sensitivity analysis assessed whether the annotation coherence of the Alzheimer packet exceeded that expected from same-size random gene sets. The case-study packet exceeded the distributions generated from 250 random sets for pathway-membership burden, total disease-association burden, and neurodegenerative-disease association burden (Figure S10A–D; Methods: Sensitivity analysis and source-dependence checks; Supplementary Note 12). This comparison indicates unusual coherence within the tested database backgrounds rather than EV specificity or function.

The case study therefore ended with a review-ready package: source-linked EV records, prioritized molecular relationships, competing interpretations, and explicit evidence gaps and verification considerations that researchers can evaluate when deciding which candidates should advance, remain uncertain, or be rejected.

## 3. Discussion

EVd3x addresses a practical bottleneck in EV research: molecular profiling can generate candidate lists faster than the supporting evidence can be reviewed. Its central contribution is not another isolated annotation layer, but a continuous, source-linked record that carries the same molecules from reported EV detection through disease, pathway, cell-context, regulatory, and interaction analyses. In the fixed Alzheimer positive control, this workflow recovered a recognizable PSEN1-centered landscape and reduced a large evidence space to a focused set of candidates, conflicts, and unresolved questions.

The value of the output lies in what it makes reviewable. EVd3x separates reported EV-cargo support from broader biological context, preserves the records behind each result, and shows where evidence layers agree or conflict. The resulting evidence-review record identifies which candidates merit attention, which interpretations should stop, and which experiments are needed before a stronger claim is made.

### 3.1 Why integrated evidence is more useful than isolated database searches

The main benefit of integration is continuity. The same resolved molecules, filters, caps, and source rows are carried through EV evidence, disease, pathway, cell-context, communication, miRNA-target, and PPI views. This reduces tool switching and hidden changes in the analyzed list while allowing agreement and disagreement to be evaluated side by side. In the case study, reported EV records, Alzheimer associations, APP-processing pathways, and miRNA-target relationships converged on a PSEN1-centered context, whereas weak cell-marker support and biologically mismatched ligand-receptor labels identified relationships that required caution or rejection.

Integration, however, is not evidence multiplication. Databases may reuse the same publications, pathway collections overlap, curated disease resources favor well-studied genes, and interaction resources may assign different molecular roles to similar relationships (Supplementary Figures S2–S3 and S7–S8; Tables S1 and S3–S7). Row counts and composite scores can therefore reflect publication density, source redundancy, or annotation breadth as well as biological relevance. EVd3x increases the breadth, continuity, and auditability of review, but it does not increase the certainty of the underlying evidence simply by combining sources.

This distinction is especially important in EV biology. A molecule can be disease-associated, expressed in a candidate source cell, included in a pathway, and connected to a receptor or target without being present in the relevant EV preparation. Even confirmed presence does not establish intravesicular localization, transfer, or activity [5–9,11,19–21,68,69]. The assistant is therefore used as a navigation and summarization layer over retrieved records rather than as an independent source of biological evidence. Table 1 and Supplementary Table S9 formalize these boundaries, while Figure S4 links each main analysis to its authoritative export.

A combined analysis should consequently end with a focused set of candidates, the evidence and uncertainty attached to each, and a defined verification requirement or stopping rule, not a single unqualified biological narrative.

### 3.2 Current limitations and verification priorities

The first limitation is inherited from the underlying evidence base. Reported EV-cargo records vary in sample type, preparation method, characterization depth, organism, disease context, and detection technology [14–17]. EV-TRACK-linked metadata improve transparency where available, but they do not resolve missing experimental detail. Most disease, pathway, expression, miRNA-target, ligand-receptor, and PPI records also originate outside the relevant EV, biofluid, cell, or disease setting. These layers therefore provide biological context only after the reported EV evidence and experimental match have been reviewed (Supplementary Note 4; Figures S2– S3; Table S3).

A second limitation concerns ranking and evaluation scope. Disease priorities, pathway enrichments, cell-context scores, communication scores, miRNA bridge scores, and PPI thresholds organize review order rather than estimate probability. The oligodendrocyte precursor and PSEN1 communication examples show why score components, molecular roles, and membrane localization must remain visible (Supplementary Notes 5–8; Figures S7–S9; Tables S4–S7). In addition, the present study evaluates one well-characterized Alzheimer query that led to a PSEN1-centered positive-control packet. The random-set analysis tests annotation coherence under the available database backgrounds, not biological function (Methods: Sensitivity analysis and source-dependence checks; Supplementary Note 12; Figure S10A–D).

Broader evaluation should therefore include blinded molecule lists, multiple cargo classes and biofluids, curated positive and negative cases, source-ablation analyses, comparison with manual expert review, and structured testing by independent EV researchers. Useful endpoints include time saved during evidence review, accuracy of claim-boundary classification, agreement on which candidates should be rejected, completeness of the verification requirements identified, and whether the exported evidence record improves research decisions.

Operational and model constraints also remain. Interactive molecule, target-scan, cell-specificity, communication, and STRING caps keep the browser responsive and have no biological meaning. The default interface accepts up to 25 input molecules per graph request; larger inputs must be divided into separate searches or processed through a separately configured workflow. Analyses should be checked for truncation flags, source dates, and database versions (Methods: Query workflow and reviewable evidence packet; Supplementary Note 3; Figure S5; Table S2). The current language model is intentionally small and local-first and may produce incomplete summaries when retrieved context is sparse or wording that overstates broad associations. Its summaries should therefore remain linked to exact exports and checked against the underlying publications (Methods: Natural-language navigation and grounding; Supplementary Notes 9–11; Figure S6; Table S8).

Safe interpretation follows three gates: establish the reported EV-associated record and preparation context; establish molecular location and relevance to the proposed cellular setting; and only then evaluate transfer or function. Failure at any gate should narrow or stop the claim rather than be offset by additional annotations. EVd3x is therefore best described as an evidence-aware platform for evidence triage, hypothesis prioritization, and identification of verification requirements, not as an EV-loading predictor, biomarker-validation engine, or substitute for expert review.

### 3.3 Building an EV evidence resource for future biological Artificial Intelligence

This first public release establishes a versioned EV knowledge base that can be expanded through expert curation and support future EV-specific Artificial Intelligence (AI) models for evidence review and hypothesis generation. However, the current knowledge base is not itself a biological training set. It’s more than 22 million records describe molecules, studies, annotations, and interactions, while the released 3,816 training examples and 424 held-out examples teach query routing and table-grounded summarization (Methods: Natural-language navigation and grounding; Supplementary Notes 9–11; Figure S6; Table S8). They do not teach a model to judge preparation quality, infer EV-mediated function, or resolve conflicting biological evidence.

A rigorous training corpus would require expert-reviewed claim–evidence units. Each unit should identify the strongest supported claim, reported detection, EV localization, selective loading, transfer, target engagement, functional response, biomarker association, or mechanism, together with the sample, preparation, characterization, assay, disease and cell context, source publication, direct or predicted support, and missing prerequisites. Hard negative and contradictory examples would be equally important, including duplicated records, context-mismatched disease associations, predicted targets without delivery evidence, poorly characterized preparations, biologically inconsistent ligand-receptor assignments, and well-studied genes whose annotation burden exceeds their EV-specific support.

An EV-specific model could then be evaluated on practical evidence-review tasks: classifying the strongest claim supported by a record set, separating independent from repeated evidence, identifying missing verification steps, and explaining why a candidate should advance, remain uncertain, or be rejected. Evaluation should use blinded expert cases, source-ablation tests, uncertainty calibration, and prospective comparison with EV researchers rather than language-quality metrics alone.

With expert-reviewed claim–evidence labels, EVd3x could evolve into an EV evidence graph, training corpus, and benchmarking environment for evidence-aware interpretation. Broader RNA classes, isolation-method and characterization filters, database snapshots, configurable null models, cohort imports, and reproducible API workflows would support that transition (Supplementary Note 13; Figure S10E–F; Table S10). Any future EV-tuned model should remain retrieval-grounded, expose the records supporting each statement, and return explicit uncertainty and stopping conditions. Its role would be to improve evidence review, not to create biological evidence.

## 4. Conclusion

EVd3x transforms a common endpoint of EV omics, a long molecular list, into a structured starting point for evidence review and hypothesis development. The same framework can also begin from a disease, pathway, cell context, or natural-language question and return a manageable, source-traceable set of molecules and relationships. By linking reported EV evidence with disease, pathway, cellular, regulatory, and interaction information within one reproducible state, EVd3x helps researchers identify coherent themes, expose conflicts, prioritize candidates, and highlight what remains unsupported or requires independent verification. The platform therefore provides an auditable bridge between high-dimensional EV data and focused, testable biological questions, while establishing a source-attributed evidence layer for future computational models. This first public release also provides a foundation for expert-curated, EV-specific AI development.

## 5. Methods

EVd3x was designed so that each displayed result could be reconstructed from a fixed analysis state and its exported source records. The methods below describe the integrated evidence architecture, identifier harmonization, query-resolution workflow, case-study construction, module-specific retrieval and ranking procedures, annotation-coherence analysis, and table-grounded natural-language navigation used to generate Figures 1–6 and the corresponding supplementary figures and tables.

### 5.1 Evidence model and analytical boundaries

The central computational unit in EVd3x is a reviewable evidence packet. Each packet records the original query, resolved molecular identifiers, visible starting molecules, inferred or expanded candidates, selected focal molecule, active filters, thresholds, processing limits, module outputs, and source-linked exports. The same packet is carried through the reported EV-cargo, disease, pathway, cell-context, ligand-receptor, miRNA-target, and protein-interaction modules so that downstream analyses do not silently substitute a different molecular set.

Evidence from different modules remains separated by role. A reported EV-cargo row records that a molecule was reported in an EV or EV-enriched resource or publication. Disease, pathway, expression, localization, cell-marker, ligand-receptor, miRNA-target, and protein-interaction records provide broader biological context but are not treated as equivalent to reported EV detection. Assistant-generated summaries are navigation outputs and do not create molecular records or evidence relationships. Table 1 and Table S9 define the interpretation supported by each evidence class and the independent verification required before a stronger biological claim can be made [14–17,23].

Unless otherwise stated, composite scores used by EVd3x are deterministic ranking heuristics fixed before the manuscript case-study analysis. They were not trained from outcome labels, optimized against biological endpoints, calibrated as probabilities, or used as inferential statistical tests. Their purpose is to organize display and review order. Component values, source rows, publication identifiers, and machine-readable exports remain the authoritative evidence underlying each rank.

No new human participants, biospecimens, patient-level datasets, animal experiments, or laboratory EV measurements were included in this study. The analyses used molecular, annotation, interaction, and study-level records available through public resources.

### 5.2 Canonical table construction and source harmonization

EVd3x was implemented as a Python web platform backed by precomputed Apache Parquet analysis tables. The manuscript release harmonized 28 public resources into 17 canonical analysis tables. Runtime description and annotation caches were excluded from the canonical-table count because they support display and lookup rather than evidence aggregation or scoring.

The 17 canonical tables and release row counts were: genes (21,763), proteins (19,343), miRNAs (2,656), molecule_summary_data (43,762), gene_annotations (19,424), ev_evidence (234,090), publication_details (3,350), miRNA_targets_scored (2,639,746), gene_pathways (404,758), gene_disease_associations (501,157), mirna_disease_associations (145), gene_expression (2,479,776), miRNA_expression (291,427), subcellular_location (13,534), cell_specificity_unified (1,587,444), ligand_receptor_pairs_full (25,779), and string_interactions (13,715,404). These counts represent a versioned analytical snapshot rather than permanent totals for the upstream resources.

Reported EV-cargo and study-metadata tables drew on ExoCarta [24], Vesiclepedia [25], SVAtlas [26], and EV-TRACK [16,17]. Molecular identity tables were built from Ensembl [27], UniProt [28], miRBase [29], and RNAcentral [30]. The miRNA-target table integrated miRTarBase [31], DIANA-TarBase [32], and TargetScan [33]. Pathway memberships were obtained from Reactome [34], KEGG [35], Gene Ontology [36], and WikiPathways [37], and disease associations were derived from DisGeNET [38]. Expression and localization information was incorporated from the Human Protein Atlas [39], RNALocate [40], miRNATissueAtlas2 [41], and miRmine [42]. Cell-context records were harmonized from the Human Protein Atlas [39], CellMarker [43], and PanglaoDB [44]. Ligand-receptor records were derived from CellPhoneDB [45,46], OmniPath Intercell [47], CellTalkDB [48], Cellinker [49], and Reactome [34], and protein-interaction records were obtained from STRING [50]. The complete source inventory, input-file classes, processing scripts, table schemas, and where-used fields are provided in Table S1 and Supplementary Note 1.

Each source-specific processor mapped records to stable canonical identifiers while retaining upstream fields needed for audit. Genes were keyed by Ensembl Gene ID with symbols retained for display. Proteins were keyed by UniProt accession and linked to their corresponding Ensembl genes. Mature miRNAs were keyed by miRBase mature accession. Publications were keyed by normalized PubMed ID, with EV-TRACK metadata joined only when an explicit EV-TRACK identifier was present in the source or publication table.

Query identifiers were normalized against a human molecular alias catalogue. Gene symbols, Ensembl identifiers, gene names, and available synonyms were mapped to canonical gene records. UniProt accessions and protein names were mapped to corresponding gene identifiers for gene-set operations while retaining protein accessions for protein-specific review. miRNA names, mature accessions, arm-specific names, and unambiguous aliases were mapped to mature miRBase records. Stem-level miRNA aliases that could refer to more than one mature arm were treated as ambiguous rather than silently assigned to one mature product.

Mixed-type identifiers were converted to comparable strings, trailing decimal artifacts were removed, and empty or null-like values were retained as missing. PubMed identifiers were restricted to canonical numeric identifiers before publication joins. Source-local Original_Evidence_ID values were not reinterpreted as EV-TRACK identifiers because they can contain database-specific or molecule-like values.

The miRNA_targets_scored processor combined experimental and predicted evidence hierarchically. miRTarBase supplied the primary experimental evidence, DIANA-TarBase added or enriched experimentally supported rows, and TargetScan added prediction-only relationships. Each unique miRNA-gene pair received a confidence score from 0 to 10 based on available assay class, TarBase support, seed-match information, and TargetScan context or probability-of-conserved-targeting fields. The original source and evidence class were retained so that experimentally supported and prediction-only relationships could be distinguished during review.

Pathway processing retained valid Ensembl identifiers, combined memberships across sources, removed duplicate source-gene-pathway records, and assigned broad pathway categories using standardized names and keyword rules. Cell-specificity processing harmonized Human Protein Atlas expression records and CellMarker and PanglaoDB marker records to common cell-type and biological-system labels. Ligand-receptor processing retained directed pair information, stimulation or inhibition annotations, source strings, and references. Source presence made a row eligible for module-specific review; it was not interpreted as evidence of biological function. The manuscript analysis used canonicalization version 2026.02.13.

### 5.3 Query workflow and reviewable evidence packet

The live EVd3x workflow supports entry through individual genes, proteins, or miRNAs; user-defined molecular collections; diseases; pathways; cell or communication contexts; and natural-language questions. Explicit molecule searches are tokenized, normalized, deduplicated, and resolved through the canonical molecule tables. Protein tokens remain available for protein details and STRING review, whereas corresponding gene identifiers are used for pathway, disease, cell-context, and target-set operations.

For generic disease-first searches, the disease label is matched to disease-association records, candidate molecules are assembled from the matched disease space, and ranked entities can be composed into a molecular query for downstream analysis. These live search routes are distinct from the frozen manuscript case study.

For the manuscript case study, we entered the exact query “early-onset Alzheimer’s disease with behavioral disturbance.” EVd3x mapped the query to familial Alzheimer disease type 3 (DisGeNET identifier C1843013), and the resulting PSEN1-centered evidence packet was recorded in Table S2 and held constant across downstream analyses. The visible starting molecules were PSEN1 mRNA, its automatically linked protein representation, and three miRNAs: hsa-miR-4496, hsa-miR-107, and hsa-miR-4677-3p. The complete candidate set also retained hsa-miR-562, hsa-miR-3064-5p, and hsa-miR-1184 as PSEN1-targeting bridge miRNAs.

The fixed gene universe contained PSEN1 together with 25 target-derived or network-context genes: APP, BACE1, MAPT, APOE, PSEN2, NOTCH1, NOTCH2, NOTCH3, NOTCH4, CTNNB1, LRRK2, IRS1, HIPK2, KDM5B, DCP1A, CBX5, CDK6, CDK8, CRKL, CSNK1G1, MAP3K2, AAK1, NFAT5, ORAI2, and CMTM4. The first-pass case-study display contained 14 ranked candidates spanning PSEN1 anchors, miRNA bridges, recommended miRNAs, and Alzheimer-relevant network neighbors. These values were loaded from the fixed Table S2 asset rather than regenerated as an unconstrained disease-discovery benchmark.

The Alzheimer analysis should therefore be interpreted as a reproducible positive-control demonstration of the EVd3x evidence workflow. It tests whether a fixed, biologically recognizable molecular state can be carried through different evidence layers without losing provenance; it does not evaluate the accuracy of unconstrained natural-language disease-to-molecule discovery.

Each downstream module read from the same recorded evidence packet. Visible starting molecules, additional bridge miRNAs, target-derived genes, and network-context genes remained separately labelled. The manuscript packet contained five visible starting molecules, 104 inferred molecules, 109 total molecular nodes, and 197 displayed relationships. The query text, molecular classes, candidate set, thresholds, expansion settings, and processing guards are reported in Table S2 and Supplementary Notes 2–3; the routing and packet-construction workflow is shown in Figure S5.

The immediate interface accepted up to 25 unique molecule tokens in a single graph request. Requests above this limit were blocked in the default release and had to be divided into separate searches or processed through a separately configured workflow. Additional retrieval limits protected the largest tables: miRNA-target scans were capped at 1,500,000 rows, cell-specificity scans at 2,000,000 rows, and STRING scans at 200,000 rows per focal protein, with an additional internal limit during graph construction. These limits are computational safeguards and have no biological meaning. Active limits, truncation status, and warnings were retained in response metadata where applicable.

Reproducible reports record the original query, entry mode, resolved identifiers, active filters, Top-N settings, processing caps, truncation flags, source snapshot or canonicalization version, and export names. Optional natural-language summarization occurs only after deterministic routing and table retrieval; molecules, disease labels, pathway memberships, and ligand-receptor or protein-interaction edges originate from the integrated tables rather than generated text.

### 5.4 EV evidence retrieval and reported-cargo provenance

Reported EV evidence was retrieved from ev_evidence by matching resolved molecular identifiers. Gene-centered views also queried linked UniProt identifiers when a cargo record was stored under a protein accession. Retrieved rows retained the canonical molecule identifier, molecule class, source database, sample type, isolation or enrichment method, detection method, original evidence identifier, and PubMed identifier where available.

Publication metadata were joined from publication_details using normalized PubMed identifiers. Available fields included title, first author, publication year, EV-TRACK identifier, sample information, isolation-protocol information, and EV-associated cell annotations. EV-TRACK identifiers were used only when explicitly present in the source metadata.

Missing sample, isolation, detection, publication, or EV-TRACK fields were retained as missing and were not converted into evidence of absence or poor study quality. Cargo rows were treated as source records rather than independent studies. When study counts were required, records were collapsed by PubMed identifier or EV-TRACK study identifier where available.

This procedure generated the reported-cargo panel in Figure 2D, the global and case-study summaries in Figures S2-S3, and the row-level EV evidence export in Table S3. Supplementary Note 4 describes the relationship between source-level, harmonized, and publication-collapsed counts.

Recommended nodes are a navigation layer over the active state. For molecule m, EVd3x records EV-evidence count C_m, source breadth S_m, and publication breadth P_m. The navigation-support term was:

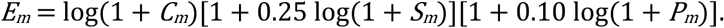

The coefficients 0.25 and 0.10 were fixed damping terms that allowed source and publication breadth to modulate ties while retaining EV-record burden as the dominant component. Disease support, pathway breadth, miRNA-target support, and graph context were displayed separately when present. Visible starting nodes and inferred nodes remained separately labelled. The recommendation score was used only for navigation and export prioritization.

### 5.5 Disease and pathway annotation

Disease analysis combined gene-disease and miRNA-disease records. Rows were grouped by disease identifier when present and otherwise by a normalized disease name. The export retained supporting entity, association type, source category, source score, publication identifier, and whether the supporting molecule belonged to the direct or expanded molecular state.

For disease group d, the live display used six normalized components: E_d, log-normalized evidence-row burden; S_d, log-normalized number of supporting molecular entities; C_d, source breadth divided by the maximum source breadth in the returned result; P_d, log-normalized publication breadth; F_d, the direct-evidence fraction; and M_d, the clipped median source-provided association score. The disease-priority score was:

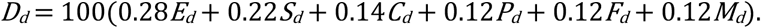

The coefficients sum to 1.00 and were fixed release defaults. The score organizes review order according to evidence burden, supporting-entity burden, source breadth, publication breadth, directness, and source score. It is not interpreted as disease probability, penetrance, diagnostic performance, causal strength, or patient-level risk. For the fixed case study, the Figure 3 disease order and selected familial Alzheimer disease type 3 record were reproduced from Table S4. Figures S7D and S8B describe source balance and direct-versus-predicted composition.

Pathway analysis used gene memberships from Reactome, KEGG, Gene Ontology, and WikiPathways. The analysis background was restricted to unique genes represented in the integrated pathway table so that enrichment was evaluated against the reachable annotation universe.

For pathway a, N was the number of unique genes in the pathway-annotated background, K_a the number of background genes assigned to pathway a, n the number of genes in the query set represented in the background, and k_a the observed query overlap. The upper-tail hypergeometric probability was:

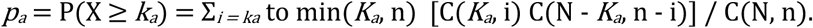

At least one overlapping gene was required when the final query set contained fewer than 50 genes, and at least two overlapping genes were required for larger sets. P values were adjusted across tested pathways using the Benjamini-Hochberg procedure [70]. The manuscript analysis used p ≤ 0.05 and q ≤ 0.10.

The complete statistically filtered result was retained in the export. A fixed manuscript display filter reduced that result to 153 visible terms across 16 categories for Figure 4, while preserving the underlying pathway memberships, overlap values, P values, and adjusted q values in Table S5. Significant terms were grouped using the harmonized pathway-category field. Category summaries retained pathway count, overlapping-gene count, and mean, median, and minimum P values. Category links were displayed only when two categories shared more than one gene and their shared-gene Jaccard index exceeded 0.1.

This procedure generated Figure 4, Figure S7A, the cutoff analysis in Figure S9, and the 2,204 pathway-membership rows in Table S5. Pathway enrichment was used to organize biological themes and review order; it was not interpreted as evidence of pathway activity.

### 5.6 Cell-context and communication prioritization

Cell-context analysis used marker and expression records from cell_specificity_unified. For each analysis state, the application constructed a final gene set containing direct gene inputs together with target-derived and network-context genes. Protein inputs were represented by their corresponding genes for this analysis.

For each gene g, expression values were standardized across cell types to obtain z_gc, the gene-specific z score in cell context c. Positive specificity was clipped and rescaled as:

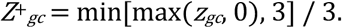

Marker information was retained separately. For expression value x_gc and marker indicator I_gc, the marker-weighted expression term was log(1 + x_gc)I_gc. Direct-seed coverage was the fraction of direct seed genes with positive specificity above 0.05 or marker support. Seed specificity was the mean positive specificity across direct seed genes, and marker support was the mean clipped marker indicator across direct seed genes.

Expanded-gene support was calculated as:

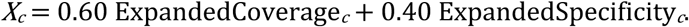

When at least one direct seed gene was present, the base context score was:

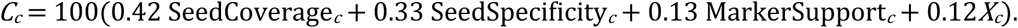

When no direct seed gene was present, the fallback score was 100(0.60X_c + 0.25 ExpandedSpecificity_c + 0.15 ExpandedCoverage_c). Communication readiness measured whether query-side ligand genes were expressed and had localization compatible with secretion or membrane association:

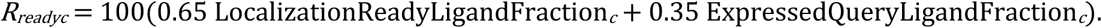

The displayed composite context score combined the base context score with bounded system relevance and communication-readiness terms:

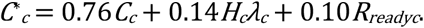

Here, H_c is the normalized system-relevance term and λ_c = min[max(2ρ, 0), 1], where ρ is the dominant-system confidence returned by the interface. The same bounded confidence factor was applied across contexts for a given query. Contexts were ordered by the composite score and then by base score, seed coverage, seed specificity, and expanded support. The component values remained visible so that a high rank driven by coverage or system weighting could be distinguished from strong cell-marker evidence. This procedure generated Figure 5A–B, Figure S8C, and the cell-context records in Table S6.

Ligand-receptor analysis intersected selected source and target cell contexts with directed pairs in ligand_receptor_pairs_full. Source-database labels, references, stimulation or inhibition annotations, and molecular direction were retained. For ligand i and receptor j, with source-cell ligand expression x_i and target-cell receptor expression y_j, the default pair-expression score was:

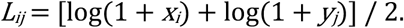

The implementation also supports product-log and geometric-log alternatives through configuration, but the manuscript used the mean-log expression score. The communication-relevance score was:

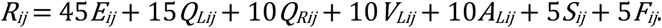

E_ij is the normalized expression score; Q^L_ij and Q^R_ij indicate direct query overlap for the ligand and receptor; V^L_ij indicates reported EV-cargo support for the ligand; A^L_ij indicates secreted or membrane annotation; S_ij is normalized curated-source breadth; and F_ij indicates agreement with the active biological-system focus. The coefficients sum to 100 and retain expression as the primary component while incorporating query overlap, reported EV support, localization compatibility, source breadth, and system context. Disease-linked pair status was retained separately and could be used for sorting or filtering, but it was not collapsed into the relevance score.

The manuscript view examined an oligodendrocyte-to-astrocyte context and hid context-only rows to focus review on records linked to the active molecular state. This procedure generated Figure 5C-D, Figure S7B, Figure S8D, and the ligand-receptor and communication records in Table S6. The scores rank candidate contexts and pairs for review; they do not infer EV cell of origin, surface display, vesicular transfer, receptor activation, or biological response.

### 5.7 miRNA-target and PPI bridge prioritization

For a set of query miRNAs, EVd3x filtered miRNA_targets_scored and grouped the retrieved rows by target gene. Experimentally supported and prediction-only source classes remained identifiable in the exported records. For target gene g, m_g was the number of distinct query miRNAs supporting the target, c-bar_g the mean confidence score, and c-max_g the maximum confidence score. The target-expansion score was:

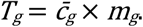

The live application supports three expansion strategies. shared_first_fill retains targets supported by at least two queried miRNAs when multiple miRNAs are present. score_rank orders all candidate targets by T_g. The legacy shared_with_fill strategy selects shared targets first and then fills remaining positions with the highest-scoring single-miRNA targets.

For the frozen manuscript case study, the authoritative final gene universe and bridge-miRNA set were loaded from Table S2. The Figure 1 network-preservation route used the legacy shared_with_fill display behavior to reproduce the submitted node and edge state. Downstream manuscript modules used the fixed Table S2 gene universe rather than re-estimating a new expansion set during each analysis.

Bridge interpretation was assigned independently from target expansion. A target was labelled direct_query_overlap when a queried miRNA targeted a queried mRNA, shared_target when at least two queried miRNAs converged on a non-query target, and single_target_context otherwise. The exported bridge score was:

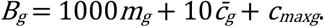

The scaling constants impose lexicographic ordering: the number of supporting query miRNAs dominates the rank, followed by mean confidence and then maximum confidence. The web graph may apply a display-only offset to direct-query-overlap relationships so that they remain visible above context-only edges. That offset was not included in the exported bridge score or interpreted as evidence strength. This procedure generated the miRNA-target components of Figure 6 and Table S7.

STRING protein-interaction rows were retrieved for the focal protein and proteins linked to genes in the active state. The manuscript view retained edges with combined_score ≥ 700, corresponding to the high-confidence STRING threshold [50]. For node-level protein-neighborhood views, rows were collapsed to canonical partner proteins before display. Reviewed UniProt entries and non-isoform-like identifiers were preferred as representatives; isoform and member identifiers were retained in the export. The maximum observed values were preserved for the STRING neighborhood, fusion, co-occurrence, co-expression, experimental, database, text-mining, and combined-score channels.

This procedure generated the PSEN1 neighborhood in Figure 2C, the protein-interaction components of Figure 6, Figure S7C, and the PPI records in Table S7. STRING edges were used as protein-network context rather than as EV-specific interaction evidence.

### 5.8 Natural-language navigation and grounding

The EVd3x assistant functions as a navigation layer over the deterministic evidence workflow. Natural-language requests are first assigned to structured intent categories, including collection lookup, molecule resolution, reported EV-evidence retrieval, disease analysis, pathway analysis, cell-context analysis, communication analysis, ligand-receptor review, PPI review, localization, export, and documentation.

After routing, the corresponding retrieval modules assemble a table-backed evidence packet. Optional language-model synthesis is invoked only after retrieval. The language model does not create the underlying molecules, source records, pathway memberships, disease associations, ligand-receptor pairs, miRNA-target relationships, PPI edges, or module-specific scores. Generated text is not stored or interpreted as an independent evidence class.

The released navigation corpus contained 3,816 training examples and 424 held-out evaluation examples across 15 intent categories. A separate 621-prompt stress suite evaluated shorthand, mixed-intent, typo-heavy, and degraded prompts. These evaluations tested intent routing, evidence retrieval, and table-grounded summarization rather than the biological accuracy of the underlying database records or the ability to adjudicate EV-specific claims.

The released adapter is available at https://huggingface.co/jonbrees/evd3x-agent-lora-qwen15b. It was fine-tuned from Qwen2.5-1.5B-Instruct using QLoRA [71–75]. Training used three epochs, a learning rate of 2 × 10^-4, maximum sequence length of 2,048, LoRA rank 16, LoRA alpha 32, dropout 0.05, batch size 4, and gradient accumulation over four steps. Table S8 records the corpus composition, model metadata, licensing, configuration, evaluation profiles, deployment constraints, and grounding policy.

The model was selected as a small, open-source, local-first navigation option. Model availability does not alter deterministic intent routing, evidence retrieval, or the source-linked exports. Generated summaries may be incomplete when retrieved context is sparse or may use wording stronger than the underlying records support. Users are therefore directed to inspect the exported rows and original publications. This procedure generated Figure 1G and is documented in Supplementary Notes 9-11, Figure S6, and Table S8.

### 5.9 Sensitivity analysis and source-dependence checks

A random-set sensitivity analysis evaluated whether the fixed Alzheimer gene packet carried more pathway and disease annotation than expected for gene sets of the same size under the available database backgrounds. The sampling universe was restricted to genes represented in both the integrated pathway and disease resources, and genes belonging to the Alzheimer case-study set were removed.

Two hundred and fifty random gene sets were sampled without replacement within each set, with each random set matched to the size of the case-study gene set. Each random set was evaluated using the same source snapshots and row-count definitions as the Alzheimer packet. Three quantities were calculated: total pathway-membership rows, total disease-association rows, and disease-association rows assigned to neurodegenerative disease categories.

The case-study values were compared with the resulting empirical distributions. These comparisons were treated as annotation-coherence and database-dependence checks. They were not biological controls and did not test EV specificity, cargo localization, transfer, pathway activity, target engagement, or function. The analysis generated Figure S10A-D and is described in Supplementary Note 12. Planned source-ablation and configurable null-model analyses are reported as future development in Supplementary Note 13 and Table S10 and were not included as current results.

### 5.10 Software implementation and reproducibility

EVd3x was implemented in Python using Flask for the web application, pandas and NumPy for data processing, PyArrow for Parquet access, and SciPy for z-score and hypergeometric calculations. Canonical evidence tables were stored as Apache Parquet files. The largest tables, including miRNA_targets_scored and string_interactions, were queried using column-projected, identifier-filtered scans with configured row limits rather than being loaded fully into memory. Smaller frequently used tables were loaded on demand and maintained in a bounded in-memory cache.

For manuscript reproduction, the analysis workflow recorded the source inventory, canonical table counts, fixed query parameters, module-specific evidence rows, scoring outputs, and interpretation boundaries used to generate Figures 1–6 and Tables S1–S10. Figure panels were formatted as vector assets only after their values and labels were checked against the corresponding evidence exports; conversion to SVG, PNG, or manuscript-layout formats did not alter the underlying records, scores, counts, or statistical results.

## 6. Data availability

The EVd3x platform is available at https://evd3x.com. Machine-readable exports supporting the Alzheimer case study are provided in the Supplementary Information and mirrored in the EVd3x companion repository at https://github.com/JonathanWeerakkody/EVd3x. The schemas, source mappings, and row counts for the 17 canonical analysis tables are documented in Table S1 and in the repository manifests.

EVd3x integrates processed records derived from multiple third-party resources. Raw upstream databases and the merged production Parquet tables are not redistributed because access and redistribution remain subject to the licensing and usage terms of the original data providers. Source manifests, table schemas, processing documentation, and manuscript-linked analysis outputs are provided in the companion repository to support reproducibility.

Read-only programmatic access is available through the external /api/v1 interface. The OpenAPI specification and API quick-reference documentation are publicly available, whereas data endpoints require a valid API key supplied through either the Authorization: Bearer <API_KEY> header or the X-API-Key header. Internal /api/* endpoints are used by the web interface and are not intended for reproducible external workflows. Programmatic analyses should report the query, analysis mode, filters, Top-N or processing-cap settings, source snapshot or date, and exported output files.

## 7. Code availability

The EVd3x companion repository is available at https://github.com/JonathanWeerakkody/EVd3x. The repository contains manuscript-linked analysis scripts, figure-and table-generation materials, table manifests, knowledge-base documentation, model-training documentation, and machine-readable supplementary outputs.

The repository represents the analysis and reproducibility release associated with this manuscript and does not contain the complete production source code or deployment configuration for the hosted EVd3x web application. Raw upstream databases and production data tables are not redistributed.

## 8. Model availability

The EVd3x navigation adapter is available at https://huggingface.co/jonbrees/evd3x-agent-lora-qwen15b. The model card provides the adapter files and associated metadata. Training-corpus composition, fine-tuning parameters, evaluation design, deployment constraints, and the table-grounding policy are summarized in Table S8.

## 9. Declaration of the use of AI-assisted technologies

During software development and manuscript preparation, the authors used Claude Code with Claude Sonnet 4.6 (Anthropic) and OpenAI Codex 5.3 to assist with code development, code review and debugging, repository organization and limited language polishing.

These tools were not used as independent sources of scientific evidence, to generate or select the underlying biological records, or to make scientific conclusions without author review. All code, analyses, interpretations, and manuscript text were reviewed, revised, and approved by the authors, who take full responsibility for the content of the work.

## 10. Acknowledgements

This work was supported by the Robert E. Leet and Clara Guthrie Patterson Trust Mentored Research Award, Bank of America, N.A., Private Bank, Trustee. We thank the researchers, curators, and software-development teams who create and maintain the public databases and knowledge resources integrated into EVd3x.

## 11. Competing interests

J.S.W. is co-founders of Eatsrock Dx Inc. and hold equity in the company. K.A.O. declares no competing interests.

## 12. Author contributions

K.A.O.: Conceptual input; extracellular-vesicle biology interpretation; manuscript writing, review, and editing; and development of the roadmap for future RNA-cargo and navigation-workflow extensions. J.S.W.: Conceptualization; methodology; software development; data curation and integration; knowledge-base construction; formal analysis; visualization; model development and fine-tuning; manuscript writing, review, and editing; project administration; and supervision. Both authors reviewed and approved the final manuscript and accept responsibility for the integrity of the work.

## Notes

https://evd3x.com/

### Summary of Updates

We substantially revised the manuscript to improve clarity, accessibility, and biological interpretation in response to the previous editorial feedback. Technical jargon has been reduced throughout, and the manuscript has been reorganized around the practical question of how extracellular vesicle researchers can move from complex molecular or disease-associated evidence to focused, experimentally testable hypotheses. The Results were rewritten as a continuous case study rather than a module-by-module description. Beginning with the complex disease query "early onset Alzheimer's disease with behavioral disturbance," the analysis demonstrates how EVd3x progressively integrates EV-associated cargo evidence with disease associations, pathways, cell context, protein interactions, and ligand-receptor relationships to converge on a manageable, PSEN1-centered set of biologically connected candidates for future experimental testing. We have clarified throughout that database associations support hypothesis generation and candidate prioritization and do not establish EV origin, delivery, or biological function. The benefits and limitations of combining heterogeneous evidence sources are now explicitly described, including evidence provenance, source dependence, conflicting or context-dependent relationships, and the need for experimental validation. The Abstract and Introduction were extensively rewritten to more clearly communicate the rationale, scale, and utility of EVd3x. The Discussion now emphasizes evidence-aware interpretation, appropriate use of the platform, experimental next steps, and the potential for the integrated, provenance-linked resource to support future EV-focused computational and AI training frameworks. The Methods and Supplementary Information were also revised for readability, reproducibility, and clearer linkage to the main Results. Figures have been retained unchanged.

### Funding

- Robert E. Leet and Clara Guthrie Patterson Trust Mentored Research Award, Bank of America, Private Bank, Trustee

## References

[1] H. Valadi, K. Ekström, A. Bossios, M. Sjöstrand, J. J. Lee, and J. O. Lötvall, “Exosome-mediated transfer of mRNAs and microRNAs is a novel mechanism of genetic exchange between cells,” Nature Cell Biology, vol. 9, no. 6, pp. 654–659, 2007, doi: 10.1038/ncb1596.

[2] G. Raposo and W. Stoorvogel, “Extracellular vesicles: Exosomes, microvesicles, and friends,” Journal of Cell Biology, vol. 200, no. 4, pp. 373–383, 2013, doi: 10.1083/jcb.201211138.

[3] S. J. Gould and G. Raposo, “As we wait: Coping with an imperfect nomenclature for extracellular vesicles,” Journal of Extracellular Vesicles, vol. 2, no. 1, p. 20389, 2013, doi: 10.3402/jev.v2i0.20389.

[4] G. van Niel, G. D’Angelo, and G. Raposo, “Shedding light on the cell biology of extracellular vesicles,” Nature Reviews Molecular Cell Biology, vol. 19, no. 4, pp. 213–228, 2018, doi: 10.1038/nrm.2017.125.

[5] J. Kowal, G. Arras, M. Colombo, et al., “Proteomic comparison defines novel markers to characterize heterogeneous populations of extracellular vesicle subtypes,” Proceedings of the National Academy of Sciences, vol. 113, no. 8, pp. E968–E977, 2016, doi: 10.1073/pnas.1521230113.

[6] M. Mathieu, L. Martin-Jaular, G. Lavieu, and C. Théry, “Specificities of secretion and uptake of exosomes and other extracellular vesicles for cell-to-cell communication,” Nature Cell Biology, vol. 21, no. 1, pp. 9–17, 2019, doi: 10.1038/s41556-018-0250-9.

[7] G. van Niel, D. R. F. Carter, A. Clayton, D. W. Lambert, G. Raposo, and P. Vader, “Challenges and directions in studying cell-cell communication by extracellular vesicles,” Nature Reviews Molecular Cell Biology, vol. 23, no. 5, pp. 369–382, 2022, doi: 10.1038/s41580-022-00460-3.

[8] A. C. Dixson, T. R. Dawson, D. Di Vizio, and A. M. Weaver, “Context-specific regulation of extracellular vesicle biogenesis and cargo selection,” Nature Reviews Molecular Cell Biology, vol. 24, no. 7, pp. 454–476, 2023, doi: 10.1038/s41580-023-00576-0.

[9] M. Kang, V. Jordan, C. Blenkiron, and L. W. Chamley, “Biodistribution of extracellular vesicles following administration into animals: A systematic review,” Journal of Extracellular Vesicles, vol. 10, no. 8, p. e12085, 2021, doi: 10.1002/jev2.12085.

[10] B. J. Benedikter, F. G. Bouwman, T. Vajen, et al., “Ultrafiltration combined with size exclusion chromatography efficiently isolates extracellular vesicles from cell culture media for compositional and functional studies,” Scientific Reports, vol. 7, no. 1, p. 15297, 2017, doi: 10.1038/s41598-017-15717-7.

[11] D. K. Jeppesen, A. M. Fenix, J. L. Franklin, et al., “Reassessment of exosome composition,” Cell, vol. 177, no. 2, pp. 428–445.e18, 2019, doi: 10.1016/j.cell.2019.02.029.

[12] Q. Zhang, D. K. Jeppesen, J. N. Higginbotham, et al., “Supermeres are functional extracellular nanoparticles replete with disease biomarkers and therapeutic targets,” Nature Cell Biology, vol. 23, no. 12, pp. 1240–1254, 2021, doi: 10.1038/s41556-021-00805-8.

[13] J. Lötvall, A. F. Hill, F. Hochberg, et al., “Minimal experimental requirements for definition of extracellular vesicles and their functions: A position statement from the International Society for Extracellular Vesicles,” Journal of Extracellular Vesicles, vol. 3, no. 1, p. 26913, 2014, doi: 10.3402/jev.v3.26913.

[14] C. Théry, K. W. Witwer, E. Aikawa, et al., “Minimal information for studies of extracellular vesicles 2018 (MISEV2018): A position statement of the international society for extracellular vesicles and update of the MISEV2014 guidelines,” Journal of Extracellular Vesicles, vol. 7, no. 1, p. 1535750, 2018, doi: 10.1080/20013078.2018.1535750.

[15] J. A. Welsh, D. C. I. Goberdhan, L. O’Driscoll, et al., “Minimal information for studies of extracellular vesicles (MISEV2023): From basic to advanced approaches,” Journal of Extracellular Vesicles, vol. 13, no. 2, p. e12404, 2024, doi: 10.1002/jev2.12404.

[16] J. Van Deun, P. Mestdagh, P. Agostinis, et al., “EV-TRACK: Transparent reporting and centralizing knowledge in extracellular vesicle research,” Nature Methods, vol. 14, no. 3, pp. 228–232, 2017, doi: 10.1038/nmeth.4185.

[17] Q. Roux, J. Van Deun, S. Dedeyne, and A. Hendrix, “The EV-TRACK summary add-on: Integration of experimental information in databases to ensure comprehensive interpretation of biological knowledge on extracellular vesicles,” Journal of Extracellular Vesicles, vol. 9, no. 1, p. 1699367, 2020, doi: 10.1080/20013078.2019.1699367.

[18] K. W. Witwer, E. I. Buzás, L. T. Bemis, et al., “Standardization of sample collection, isolation and analysis methods in extracellular vesicle research,” Journal of Extracellular Vesicles, vol. 2, no. 1, p. 20360, 2013, doi: 10.3402/jev.v2i0.20360.

[19] A. F. Hill, D. M. Pegtel, U. Lambertz, et al., “ISEV position paper: Extracellular vesicle RNA analysis and bioinformatics,” Journal of Extracellular Vesicles, vol. 2, no. 1, p. 22859, 2013, doi: 10.3402/jev.v2i0.22859.

[20] B. Mateescu, E. J. K. Kowal, B. W. M. van Balkom, et al., “Obstacles and opportunities in the functional analysis of extracellular vesicle RNA–an ISEV position paper,” Journal of Extracellular Vesicles, vol. 6, no. 1, p. 1286095, 2017, doi: 10.1080/20013078.2017.1286095.

[21] R. T. Miceli, T.-Y. Chen, Y. Nose, et al., “Extracellular vesicles, RNA sequencing, and bioinformatic analyses: Challenges, solutions, and recommendations,” Journal of Extracellular Vesicles, vol. 13, no. 12, p. e70005, 2024, doi: 10.1002/jev2.70005.

[22] U. Erdbrügger, C. J. Blijdorp, I. V. Bijnsdorp, et al., “Urinary extracellular vesicles: A position paper by the Urine Task Force of the International Society for Extracellular Vesicles,” Journal of Extracellular Vesicles, vol. 10, no. 7, p. e12093, 2021, doi: 10.1002/jev2.12093.

[23] R. Crescitelli, J. M. Falcón-Pérez, A. Hendrix, et al., “Reproducibility of extracellular vesicle research,” Journal of Extracellular Vesicles, vol. 14, no. 1, p. e70036, 2025, doi: 10.1002/jev2.70036.

[24] S. Mathivanan and R. J. Simpson, “ExoCarta: A compendium of exosomal proteins and RNA,” Proteomics, vol. 9, no. 21, pp. 4997–5000, 2009, doi: 10.1002/pmic.200900351.

[25] S. V. Chitti, S. Gummadi, T. Kang, et al., “Vesiclepedia 2024: An extracellular vesicles and extracellular particles repository,” Nucleic Acids Research, vol. 52, no. D1, pp. D1694–D1698, 2024, doi: 10.1093/nar/gkad1007.

[26] Z. Wei, N. Zhou, M. Jing, et al., “SVAtlas: A comprehensive single extracellular vesicle omics resource,” Nucleic Acids Research, vol. 54, no. D1, pp. D1807–D1816, 2026, doi: 10.1093/nar/gkaf1189.

[27] P. W. Harrison, M. R. Amode, O. Austine-Orimoloye, et al., “Ensembl 2024,” Nucleic Acids Research, vol. 52, no. D1, pp. D891–D899, 2024, doi: 10.1093/nar/gkad1049.

[28] The UniProt Consortium, “UniProt: The Universal Protein Knowledgebase in 2023,” Nucleic Acids Research, vol. 51, no. D1, pp. D523–D531, 2023, doi: 10.1093/nar/gkac1052.

[29] A. Kozomara and S. Griffiths-Jones, “miRBase: Annotating high confidence microRNAs using deep sequencing data,” Nucleic Acids Research, vol. 42, no. D1, pp. D68–D73, 2014, doi: 10.1093/nar/gkt1181.

[30] The RNAcentral Consortium, “RNAcentral 2021: Secondary structure integration, improved sequence search and new member databases,” Nucleic Acids Research, vol. 49, no. D1, pp. D212–D220, 2021, doi: 10.1093/nar/gkaa921.

[31] H.-Y. Huang, Y.-C.-D. Lin, S. Cui, et al., “miRTarBase update 2022: An informative resource for experimentally validated miRNA-target interactions,” Nucleic Acids Research, vol. 50, no. D1, pp. D222–D230, 2022, doi: 10.1093/nar/gkab1079.

[32] D. Karagkouni, M. D. Paraskevopoulou, S. Chatzopoulos, et al., “DIANA-TarBase v8: A decade-long collection of experimentally supported miRNA-gene interactions,” Nucleic Acids Research, vol. 46, no. D1, pp. D239–D245, 2018, doi: 10.1093/nar/gkx1141.

[33] V. Agarwal, G. W. Bell, J.-W. Nam, and D. P. Bartel, “Predicting effective microRNA target sites in mammalian mRNAs,” eLife, vol. 4, p. e05005, 2015, doi: 10.7554/eLife.05005.

[34] M. Gillespie, B. Jassal, R. Stephan, et al., “The Reactome pathway knowledgebase 2022,” Nucleic Acids Research, vol. 50, no. D1, pp. D687–D692, 2022, doi: 10.1093/nar/gkab1028.

[35] M. Kanehisa, M. Furumichi, Y. Sato, M. Kawashima, and M. Ishiguro-Watanabe, “KEGG for taxonomy-based analysis of pathways and genomes,” Nucleic Acids Research, vol. 51, no. D1, pp. D587–D592, 2023, doi: 10.1093/nar/gkac963.

[36] The Gene Ontology Consortium, “The Gene Ontology knowledgebase in 2023,” Genetics, vol. 224, no. 1, p. iyad031, 2023, doi: 10.1093/genetics/iyad031.

[37] M. Martens, A. Ammar, A. Riutta, et al., “WikiPathways: Connecting communities,” Nucleic Acids Research, vol. 49, no. D1, pp. D613–D621, 2021, doi: 10.1093/nar/gkaa1024.

[38] J. Piñero, J. M. Ramírez-Anguita, J. Saüch-Pitarch, et al., “The DisGeNET knowledge platform for disease genomics: 2019 update,” Nucleic Acids Research, vol. 48, no. D1, pp. D845–D855, 2020, doi: 10.1093/nar/gkz1021.

[39] P. J. Thul and C. Lindskog, “The Human Protein Atlas: A spatial map of the human proteome,” Protein Science, vol. 27, no. 1, pp. 233–244, 2018, doi: 10.1002/pro.3307.

[40] T. Cui, Y. Dou, P. Tan, et al., “RNALocate v2.0: An updated resource for RNA subcellular localization with increased coverage and annotation,” Nucleic Acids Research, vol. 50, no. D1, pp. D333–D339, 2022, doi: 10.1093/nar/gkab825.

[41] A. Keller, L. Gröger, T. Tschernig, et al., “miRNATissueAtlas2: An update to the human miRNA tissue atlas,” Nucleic Acids Research, vol. 50, no. D1, pp. D211–D221, 2022, doi: 10.1093/nar/gkab808.

[42] B. Panwar, G. S. Omenn, and Y. Guan, “miRmine: A database of human miRNA expression profiles,” Bioinformatics, vol. 33, no. 10, pp. 1554–1560, 2017, doi: 10.1093/bioinformatics/btx019.

[43] X. Zhang, Y. Lan, J. Xu, et al., “CellMarker: A manually curated resource of cell markers in human and mouse,” Nucleic Acids Research, vol. 47, no. D1, pp. D721–D728, 2019, doi: 10.1093/nar/gky900.

[44] O. Franzén, L.-M. Gan, and J. L. M. Björkegren, “PanglaoDB: A web server for exploration of mouse and human single-cell RNA sequencing data,” Database, vol. 2019, p. baz046, 2019, doi: 10.1093/database/baz046.

[45] R. Vento-Tormo, M. Efremova, R. A. Botting, et al., “Single-cell reconstruction of the early maternal-fetal interface in humans,” Nature, vol. 563, no. 7731, pp. 347–353, 2018, doi: 10.1038/s41586-018-0698-6.

[46] M. Efremova, M. Vento-Tormo, S. A. Teichmann, and R. Vento-Tormo, “CellPhoneDB: Inferring cell-cell communication from combined expression of multi-subunit ligand-receptor complexes,” Nature Protocols, vol. 15, no. 4, pp. 1484–1506, 2020, doi: 10.1038/s41596-020-0292-x.

[47] D. Türei, A. Valdeolivas, L. Gul, et al., “Integrated intra- and intercellular signaling knowledge for multicellular omics analysis,” Molecular Systems Biology, vol. 17, no. 3, p. e9923, 2021, doi: 10.15252/msb.20209923.

[48] X. Shao, J. Liao, C. Li, X. Lu, J. Cheng, and X. Fan, “CellTalkDB: A manually curated database of ligand-receptor interactions in humans and mice,” Briefings in Bioinformatics, vol. 22, no. 4, p. bbaa269, 2021, doi: 10.1093/bib/bbaa269.

[49] Y. Zhang, T. Liu, J. Wang, et al., “Cellinker: A platform of ligand-receptor interactions for intercellular communication analysis,” Bioinformatics, vol. 37, no. 14, pp. 2025–2032, 2021, doi: 10.1093/bioinformatics/btab036.

[50] D. Szklarczyk, R. Kirsch, M. Koutrouli, et al., “The STRING database in 2023: Protein-protein association networks and functional enrichment analyses for any sequenced genome of interest,” Nucleic Acids Research, vol. 51, no. D1, pp. D638–D646, 2023, doi: 10.1093/nar/gkac1000.

[51] A. Goate, M.-C. Chartier-Harlin, M. Mullan, et al., “Segregation of a missense mutation in the amyloid precursor protein gene with familial Alzheimer’s disease,” Nature, vol. 349, no. 6311, pp. 704–706, 1991, doi: 10.1038/349704a0.

[52] J. A. Hardy and G. A. Higgins, “Alzheimer’s disease: The amyloid cascade hypothesis,” Science, vol. 256, no. 5054, pp. 184–185, 1992, doi: 10.1126/science.1566067.

[53] R. Sherrington, E. I. Rogaev, Y. Liang, E. A. Rogaeva, G. Levesque, et al., “Cloning of a gene bearing missense mutations in early-onset familial Alzheimer’s disease,” Nature, vol. 375, no. 6534, pp. 754–760, 1995, doi: 10.1038/375754a0.

[54] E. H. Corder, A. M. Saunders, W. J. Strittmatter, et al., “Gene dose of apolipoprotein E type 4 allele and the risk of Alzheimer’s disease in late onset families,” Science, vol. 261, no. 5123, pp. 921–923, 1993, doi: 10.1126/science.8346443.

[55] B. De Strooper, P. Saftig, K. Craessaerts, H. Vanderstichele, et al., “Deficiency of presenilin-1 inhibits the normal cleavage of amyloid precursor protein,” Nature, vol. 391, no. 6665, pp. 387–390, 1998, doi: 10.1038/34910.

[56] W. X. Wang, B. J. Rajeev, A. J. Stromberg, N. Ren, et al., “The expression of microRNA miR-107 decreases early in Alzheimer’s disease and may accelerate disease progression through regulation of beta-site amyloid precursor protein-cleaving enzyme 1,” The Journal of Neuroscience, vol. 28, no. 5, pp. 1213–1223, 2008, doi: 10.1523/JNEUROSCI.5065-07.2008.

[57] C. M. Karch and A. M. Goate, “Alzheimer’s disease risk genes and mechanisms of disease pathogenesis,” Biological Psychiatry, vol. 77, no. 1, pp. 43–51, 2015, doi: 10.1016/j.biopsych.2014.05.006.

[58] H. Mathys et al., “Single-cell transcriptomic analysis of Alzheimer’s disease,” Nature, vol. 570, no. 7761, pp. 332–337, 2019, doi: 10.1038/s41586-019-1195-2.

[59] A. Grubman et al., “A single-cell atlas of entorhinal cortex from individuals with Alzheimer’s disease reveals cell-type-specific gene expression regulation,” Nature Neuroscience, vol. 22, no. 12, pp. 2087–2097, 2019, doi: 10.1038/s41593-019-0539-4.

[60] D. M. Walsh et al., “Naturally secreted oligomers of amyloid beta protein potently inhibit hippocampal long-term potentiation in vivo,” Nature, vol. 416, no. 6880, pp. 535–539, 2002, doi: 10.1038/416535a.

[61] J. P. Cleary et al., “Natural oligomers of the amyloid-beta protein specifically disrupt cognitive function,” Nature Neuroscience, vol. 8, no. 1, pp. 79–84, 2005, doi: 10.1038/nn1372.

[62] T. Ma et al., “Suppression of eIF2alpha kinases alleviates alzheimer’s disease-related plasticity and memory deficits,” Nature Neuroscience, vol. 16, no. 9, pp. 1299–1305, 2013, doi: 10.1038/nn.3486.

[63] L. Rajendran, M. Honsho, T. R. Zahn, P. Keller, et al., “Alzheimer’s disease beta-amyloid peptides are released in association with exosomes,” Proceedings of the National Academy of Sciences of the United States of America, vol. 103, no. 30, pp. 11172–11177, 2006, doi: 10.1073/pnas.0603838103.

[64] K. Yuyama, H. Sun, S. Mitsutake, and Y. Igarashi, “A potential function for neuronal exosomes: Sequestering intracerebral amyloid-beta peptide,” FEBS Letters, vol. 589, no. 1, pp. 84–88, 2015, doi: 10.1016/j.febslet.2014.11.027.

[65] M. S. Fiandaca et al., “Identification of preclinical Alzheimer’s disease by a profile of pathogenic proteins in neurally derived blood exosomes: A case-control study,” Alzheimer’s & Dementia, vol. 11, no. 6, pp. 600–607.e1, 2015, doi: 10.1016/j.jalz.2014.06.008.

[66] E. J. Goetzl et al., “Altered lysosomal proteins in neural-derived plasma exosomes in preclinical Alzheimer disease,” Neurology, vol. 85, no. 1, pp. 40–47, 2015, doi: 10.1212/WNL.0000000000001702.

[67] E. J. Goetzl et al., “Decreased synaptic proteins in neuronal exosomes of frontotemporal dementia and Alzheimer’s disease,” FASEB Journal, vol. 30, no. 12, pp. 4141–4148, 2016, doi: 10.1096/fj.201600816R.

[68] R. R. Mizenko, M. Feaver, B. T. Bozkurt, et al., “A critical systematic review of extracellular vesicle clinical trials,” Journal of Extracellular Vesicles, vol. 13, no. 10, p. e12510, 2024, doi: 10.1002/jev2.12510.

[69] C. Fusco, G. De Rosa, I. Spatocco, et al., “Extracellular vesicles as human therapeutics: A scoping review of the literature,” Journal of Extracellular Vesicles, vol. 13, no. 5, p. e12433, 2024, doi: 10.1002/jev2.12433.

[70] Y. Benjamini and Y. Hochberg, “Controlling the false discovery rate: A practical and powerful approach to multiple testing,” Journal of the Royal Statistical Society: Series B (Methodological), vol. 57, no. 1, pp. 289–300, 1995, doi: 10.1111/j.2517-6161.1995.tb02031.x.

[71] J. Bai, et al., “Qwen technical report,” arXiv preprint arXiv:2309.16609, 2023, doi: 10.48550/arXiv.2309.16609.

[72] Qwen Team, “Qwen2.5 technical report,” arXiv preprint arXiv:2412.15115, 2024, doi: 10.48550/arXiv.2412.15115.

[73] E. J. Hu, et al., “LoRA: Low-rank adaptation of large language models,” arXiv preprint arXiv:2106.09685, 2021, doi: 10.48550/arXiv.2106.09685.

[74] T. Dettmers, A. Pagnoni, A. Holtzman, and L. Zettlemoyer, “QLoRA: Efficient finetuning of quantized LLMs,” arXiv preprint arXiv:2305.14314, 2023, doi: 10.48550/arXiv.2305.14314.

[75] T. Wolf, L. Debut, V. Sanh, et al., “Transformers: State-of-the-art natural language processing,” Proceedings of the 2020 Conference on Empirical Methods in Natural Language Processing: System Demonstrations, pp. 38–45, 2020, doi: 10.18653/v1/2020.emnlp-demos.6.

