## Supplemental Information for "EVd3x: A Bioinformatics Platform for Evidence-aware Interpretation of EV Cargo"

### 1 Supplementary overview: the evidence trail behind the EVd3x case study

The Supplementary Information provides the complete audit trail for the PSEN1-centered Alzheimer case study presented in the main manuscript. It connects each main figure to the public resources, query settings, source rows, scoring components, sensitivity analyses, and machine-readable exports used to generate the displayed results.

EVd3x supports two complementary entry routes. Researchers can begin with an experimental cargo list and add biological context, or they can begin with a disease, pathway, cell-context, or natural-language question and use the connected evidence layers to identify a manageable set of candidate molecules and relationships. In the worked example, the query “early-onset Alzheimer’s disease with behavioral disturbance” led to a focused PSEN1-centered evidence packet.

The PSEN1-centered packet was recorded in Table S2 and held constant as it was followed step by step through reported EV cargo, disease and pathway annotations, cell context, ligand–receptor relationships, miRNA–target evidence, and protein-interaction networks. The same visible starting molecules, inferred candidates, filters, thresholds, and source rows were retained across modules so that each result could be traced to the evidence that produced it.

The supplementary analyses organize information that researchers can use to formulate hypotheses and develop appropriate experimental plans; they do not establish EV loading, molecular localization, transfer, target engagement, or biological function. The interpretation boundaries follow MISEV2014, MISEV2018, MISEV2023, EV-TRACK, EV-RNA, and EV reproducibility recommendations [1–7]. Candidates are therefore retained, questioned, or prioritized for independent verification rather than presented as experimentally validated conclusions.

Figures S1–S4 establish the source architecture, cargo-database and EV-TRACK coverage, case-study EV-evidence provenance, and the mapping between main figures and machine-readable exports. Figures S5–S10 document query routing, assistant grounding, source and threshold dependence, cross-module annotation summaries, pathway cutoff behavior, sensitivity analyses, and planned infrastructure. Tables S1–S10 contain the complete source inventory, fixed-query parameters, row-level evidence, score components, interpretation boundaries, and development roadmap.

### 2 Supplementary Note 1: Source roles and EV-record provenance

EVd3x integrates resources that play distinct biological and computational roles. A molecule may appear as reported EV cargo, a disease-associated gene, a pathway member, a cell marker, a miRNA target, a ligand-receptor participant, or a protein-interaction neighbor. These roles remain separate within the shared evidence packet so that broader biological context does not become equivalent to direct EV evidence. The source roles support the Introduction, Figure 1, Methods: Canonical table construction and source harmonization, Figure S1, and Table S1.

| Layer | Example sources | Canonical key | Role in the main manuscript |
|---|---|---|---|
| **Identity mapping** | Ensembl [8], UniProt [9], miRBase [10], RNAcentral [11] | Ensembl Gene ID, UniProt accession, miRBase mature accession | Query resolution and identifier checks |

| Layer | Example sources | Canonical key | Role in the main manuscript |
|---|---|---|---|
| **EV evidence** | ExoCarta [12], Vesiclepedia [13], SVAtlas [14], EV-TRACK [4,5] | Molecule ID, PubMed ID | Figures 1–2: reported EV evidence and provenance |
| **miRNA targets** | miRTarBase [15], DIANA-TarBase [16], TargetScan [17] | miRBase mature accession, Ensembl Gene ID | Figure 6: candidate regulatory relationships requiring verification |
| **Pathways** | Reactome [18], KEGG [19], Gene Ontology [20], WikiPathways [21] | Ensembl Gene ID | Figure 4: biological themes that can guide subsequent testing |
| **Disease associations** | DisGeNET [22] (including source categories such as ClinVar, OMIM, and Orphanet) | Disease ID, Gene ID, miRNA ID | Figure 3: disease-context coherence and literature review |
| **Expression and localization** | Human Protein Atlas [23], RNALocate [24], miRNATissueAtlas2 [25], miRmine [26] | Gene ID, miRNA ID | Figures 2 and 5: localization and cellular context |
| **Cell specificity** | CellMarker [27], PanglaoDB [28], Human Protein Atlas [23] | Gene ID, cell label | Figure 5: plausible cell settings for review |
| **Communication and PPI** | CellPhoneDB [29,30], OmniPath Intercell [31], CellTalkDB [32], Cellinker [33], Reactome LRdb [18], STRING [34] | Gene symbol, STRING protein ID | Figures 5–6: communication and interaction checks |

The current counts represent a versioned snapshot rather than a complete catalogue of EV cargo or biological knowledge. Some reported cargo may not yet be indexed by the integrated repositories, and EV-TRACK and other source databases continue to expand. EVd3x is designed to incorporate updated source snapshots while retaining database versions, processing provenance, and the fixed exports required to reproduce earlier analyses.

#### 3 Supplementary Note 2: Building the PSEN1-centered Alzheimer evidence packet

This note records how the complex natural-language disease query led to the visible starting molecules, inferred candidates, thresholds, and processing settings used throughout Figures 1–6. The packet was recorded in Table S2 and held constant across downstream analyses.

The canonical tables cover identity mapping, molecule summaries, EV evidence, publication metadata, disease associations, pathway memberships, expression, localization, cell specificity, ligand-receptor pairs, miRNA targets, and STRING interactions. Runtime description caches support lookup and display but are not counted as evidence layers.

Source-specific processors standardize identifiers while retaining upstream fields needed for audit. Identity records are anchored to Ensembl, UniProt, miRBase, and RNAcentral; reported EV cargo to ExoCarta, Vesiclepedia, SVAtlas, and EV-TRACK-linked metadata; and broader context to target, pathway, disease, expression, localization, cell, communication, and PPI resources.

Table S2 records the complete Alzheimer evidence packet. The visible starting molecules were PSEN1 mRNA, its corresponding protein representation, hsa-miR-4496, hsa-miR-107, and hsa-miR-4677-3p. The retained bridge set also included hsa-miR-562, hsa-miR-3064-5p, and hsa-miR-1184 because the integrated miRNA–target table linked them to PSEN1. Expanded genes provided APP-processing, presenilin/Notch, kinase, and network-neighbor context. Visible starting molecules, bridge miRNAs, inferred candidates, and broader network context remained separately labelled so that the basis for each inclusion could be reviewed. This recorded packet provides a reproducible positive-control demonstration; it is not an evaluation of unconstrained natural-language disease-to-molecule discovery.

### 4 Supplementary Note 3: Query entry, expansion, and processing controls

EVd3x supports molecule-first, collection-based, and biology-first entry routes, including disease, pathway, cell-context, communication, and natural-language queries. Generic disease-first searches assemble candidates from matched disease-association records; the manuscript case study instead follows the recorded PSEN1-centered packet in Table S2. Figure S5 shows how explicit molecule searches and natural-language queries enter different routing steps before downstream table-backed review.

Explicit search parses genes, proteins, and miRNAs. Protein identifiers remain available for molecule details and PPI review, while linked genes support gene-set operations. The default interface accepts up to 25 unique molecule tokens per graph request; requests above this limit are blocked and must be divided into separate searches or processed through a separately configured workflow. Target-scan, cell-specificity, communication, and STRING limits keep the application responsive; all are processing controls with no biological meaning.

Expansion changes the active evidence packet and should therefore be reported with the query text, entry mode, filters, Top-N settings, caps, truncation flags, source snapshot or date, and export names. Figure S5 and Table S2 record these values for the Alzheimer run.

### 5 Supplementary Note 4: Source-linked EV cargo and publication evidence

The EV evidence layer preserves the sample, preparation, detection, publication, and source-database context needed to interpret each reported cargo record.

Table S3 contains 647 EV-evidence rows for the complete Alzheimer packet, and Figures S2–S3 summarize database coverage, PubMed linkage, EV-TRACK linkage, sample context, and molecule class. Rows should be collapsed by PubMed or EV-TRACK study identifier before record counts are interpreted as study counts. Figure S2A reports source-level pre-harmonization tallies, whereas Table S1 reports the deduplicated canonical-table count; the processing stage should therefore be named whenever these values are cited. Neither view is a complete catalogue of all EV cargo reported in the literature, because source databases and EV-TRACK continue to grow.

#### 6 Supplementary Note 5: Disease and pathway evidence supporting the case-study narrative

Disease and pathway resources are used to place the recorded PSEN1-centered packet within the expected Alzheimer context and to organize its annotation breadth into themes that researchers may consider when developing subsequent studies.

The Alzheimer export contains 4,053 disease rows. The disease-priority score ranks review order using row burden, supporting-entity burden, source breadth, publication breadth, direct-evidence fraction, and source score. The raw rows in Table S4 remain authoritative, and a high rank should not be interpreted as diagnostic probability, biomarker performance, or causal evidence.

Figures S7D and S8B show source imbalance and direct-versus-predicted evidence balance. These panels make clear that extensive curation and repeated literature can increase a familiar gene's rank. Their purpose is to explain the score and expose literature-density bias, not to strengthen the EV-specific claim.

Table S5 contains 2,204 pathway-membership rows from Reactome, KEGG, Gene Ontology, and WikiPathways. Recurrent Alzheimer, Notch/gamma-secretase, phosphorylation, neurotrophin, synaptic, and vesicle-transport themes reduce the active list to candidate biological branches for researcher review and hypothesis development. They do not show pathway activity in an EV-producing or recipient cell.

Different pathway counts refer to distinct analysis stages: the broad packet footprint, the manuscript-visible filtered terms, the full significant-term set, and the membership rows exported in Table S5. Figure S9 records cutoff behavior based on hypergeometric testing with Benjamini–Hochberg correction [35], keeping the denominator and filtering choices visible.

#### 7 Supplementary Note 6: Cell-context and communication candidates for further verification

Cell-context and communication modules place molecular evidence within plausible cellular settings by combining marker, expression, localization, and interaction records. These outputs identify contexts and relationships for review; they do not establish EV cell of origin, recipient-cell exposure, or communication.

The cell module builds a final gene set from direct inputs plus expanded support genes, scans cell_specificity_unified, computes per-gene specificity across cell labels, and keeps marker support separate from expression support. Displayed components include seed coverage, seed specificity, marker support, expanded support, system relevance, and communication readiness. Component values should be interpreted before the composite rank.

Communication analysis intersects selected source and target cell contexts with curated ligand-receptor pairs. The Figure 5 score combines expression, query-side ligand or receptor status, reported EV-ligand support, secreted or membrane annotation, source breadth, and active system focus. Table S6 retains the component evidence so that biologically mismatched molecular roles can be questioned or rejected before researchers consider these relationships for experimental follow-up.

#### 8 Supplementary Note 7: miRNA-target relationships requiring regulatory verification

The miRNA-target layer connects queried miRNAs to candidate mRNAs and retains the source and evidence class for every relationship.

The miRNA_targets_scored table integrates miRTarBase, TarBase, and TargetScan with confidence fields. Target expansion prioritizes support from multiple queried miRNAs before high-confidence single-miRNA targets, and bridge ranking separates direct query overlap, shared targets, and single-target context.

For the Alzheimer packet, six miRNAs retained in the case-study packet have target support for PSEN1, and several shared-target rows connect these miRNAs to kinase, chromatin, or network-context genes. Table S7 stores the evidence fields used by the Figure 6 bridge view. These records identify relationships that may warrant qPCR, protein measurement, perturbation, or reporter assays only after EV localization and transfer have been independently addressed.

#### 9 Supplementary Note 8: Ligand-receptor and protein-network relationships for candidate triage

Ligand-receptor and protein-interaction records are used to compare candidate molecular relationships and identify those that warrant independent verification or closer review.

Node-level STRING rows are collapsed to canonical protein partners while preserving member identifiers and maximum evidence-channel scores. Figure S7 records the high-confidence STRING threshold and ligand-receptor source mix used in the manuscript view. Biologically mismatched pairs remain useful as rejection or clarification examples because they show where integration must stop rather than produce a mechanism.

#### 10 Supplementary Note 9: Table-grounded assistant routing and evidence retrieval

The assistant functions as a navigation layer over retrieved EVd3x records, allowing natural-language questions to reach the relevant evidence tables and exports.

Table S8 and Figure S6 record the intent categories, training-corpus layers, QLoRA configuration, held-out validation set, and post-training stress prompts. These evaluations describe routing, retrieval, and summarization behavior rather than biological accuracy, EV preparation quality, or EV function.

#### 11 Supplementary Note 10: Navigation training data and a future EV evidence corpus

The current training set teaches routing and evidence summarization across EVd3x modules and provides the starting point for a larger expert-labeled EV evidence resource.

The released package contains 3,816 training examples and 424 held-out validation examples across 15 intent categories. A separate 621-prompt stress suite audits shorthand, mixed-intent, typo-heavy, and degraded inputs. These sets test whether the assistant reaches and summarizes the correct evidence tables; they do not test whether it can adjudicate an EV-specific claim.

A future biological training corpus would require expert-reviewed claim-evidence units. Each unit should label the strongest supported claim - reported detection, location inside or on an EV, selective loading, transfer, target engagement, functional response, biomarker association, or mechanism - together with the sample, preparation method, EV characterization, assay, disease and cell context, source publication, direct or predicted support, and missing prerequisites.

The corpus should also contain hard negative and contradictory examples, including duplicated records, context-mismatched disease associations, predicted targets without delivery evidence, poorly characterized EV preparations, and ligand-receptor assignments that conflict with known molecular roles. These examples are needed to prevent a future model from equating connectivity, row count, or literature density with experimental sufficiency.

#### 12 Supplementary Note 11: Model release, reproducibility, and deployment

The released adapter and model metadata make the navigation layer reproducible across local or resource-limited deployments.

The adapter is available at https://huggingface.co/jonbrees/evd3x-agent-lora-qwen15b. It uses LoRA [36] and QLoRA [37] to fine-tune Qwen2.5-1.5B-Instruct, from the Qwen model family [38,39]. Table S8 records the configuration, licensing, deployment constraints, and grounding policy.

Because the current model is intentionally small and local-first, generated summaries can be incomplete or overly fluent when retrieved context is sparse or dominated by broad annotations. Users should cite and inspect exported rows and source publications rather than treating assistant text as evidence.

#### 13 Supplementary Note 12: Sensitivity to source coverage and database background

The sensitivity analysis evaluates whether the Alzheimer packet shows unusual annotation coherence relative to same-size random gene sets under the available database backgrounds.

The sensitivity analysis samples 250 same-size gene sets from genes represented in both pathway and disease resources, excluding the case genes. The null panels compare pathway-row burden, total disease-row burden, and neurodegenerative-disease-row burden with the case-study packet.

Table S9 records what each evidence layer retains, what it cannot establish, and which experiment is required before a stronger claim can be made. The random-set comparisons are database-dependence checks and do not replace EV characterization, molecular perturbation, transfer, or functional experiments.

#### 14 Supplementary Note 13: Roadmap for an evidence-aware EV research infrastructure

This first public release establishes a versioned EV knowledge base that can be expanded through expert curation and support future EV-specific AI models for evidence review and hypothesis generation. Planned extensions include broader RNA classes, source-ablation analyses, configurable null models, database snapshots, cohort-level imports, and expert-reviewed claim–evidence labels. These are roadmap items, not capabilities evaluated in the current case study.

### 15 Supplementary Figures

The supplementary figures follow the order in which their evidence is introduced in the main manuscript. They document data provenance, query construction, analysis settings, source dependence, and interpretation boundaries without treating annotation density as experimental validation.

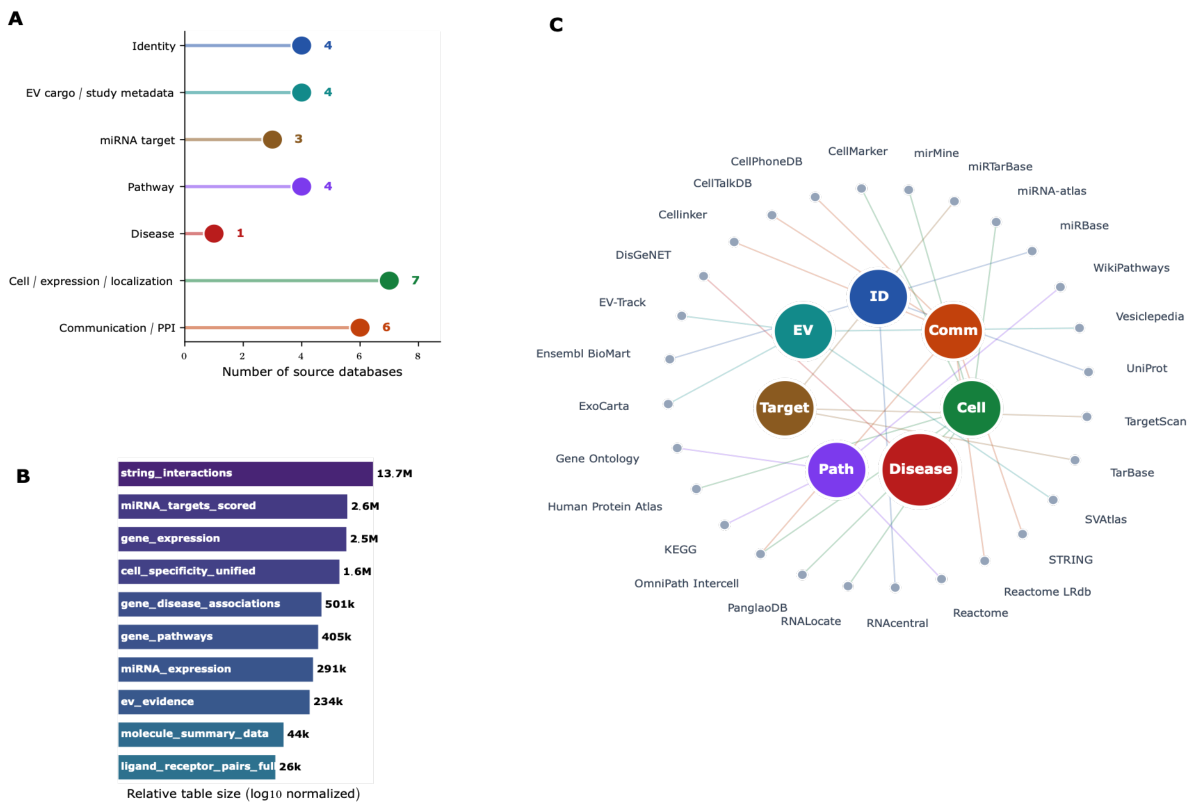

**Figure S1.** Source-role map for the EVd3x evidence-aware workflow. **A)** Source families are grouped by evidence role. **B)** Canonical analysis-table sizes are shown on a log scale. **C)** A bipartite provenance map connects source databases to evidence roles. This figure supports Figure 1 and Methods: Canonical table construction and source harmonization; the complete inventory is in Table S1. Layout position and source count describe organization and coverage, not evidence strength.

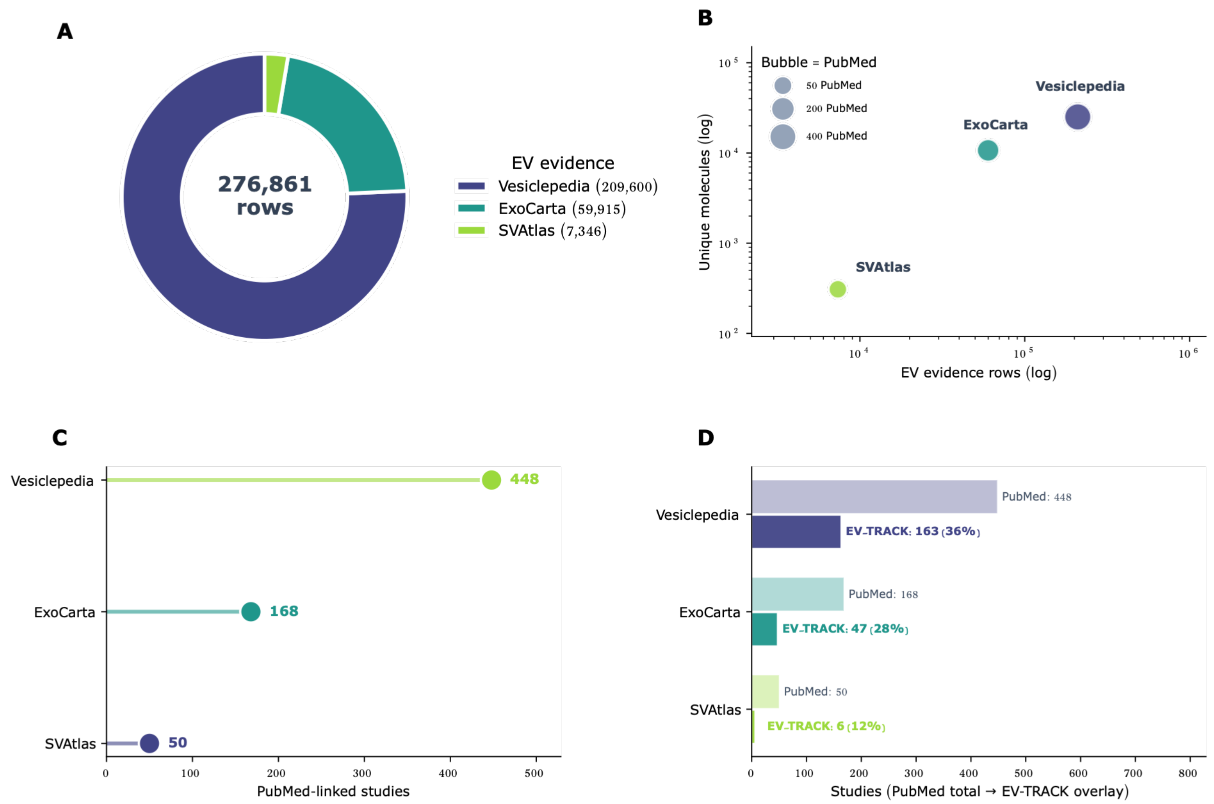

**Figure S2.** Cargo-database support, publication linkage, and EV-TRACK-linked metadata for reported EV evidence. **A)** Source-level row burden is shown before harmonization. **B)** Molecule coverage is compared across sources, with bubble size indicating PubMed linkage. **C)** PubMed-linked studies are counted by source. **D)** EV-TRACK-linked metadata are overlaid on PubMed-linked studies. The source-level tally and the harmonized canonical-table count describe different processing stages. This figure supports Results Section 2.2, Figure 2D, and Methods: EV evidence retrieval and reported-cargo provenance; the case-packet rows are in Table S3. Row counts are not study counts unless collapsed by PubMed or EV-TRACK study identifier.

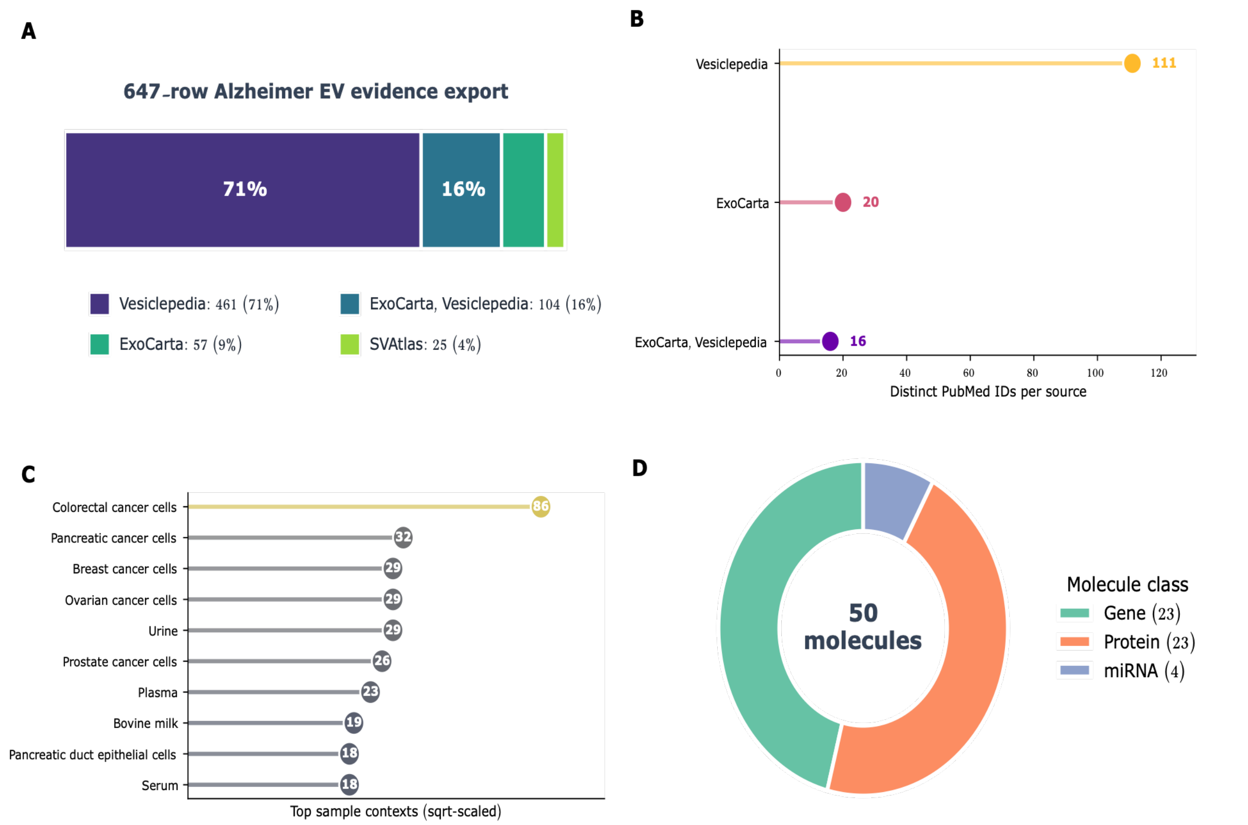

**Figure S3.** Source-attributed EV evidence for the Alzheimer case-study packet. **A)** The 647-row export is summarized by source composition. **B)** Distinct PubMed identifiers are counted by source. **C)** The leading sample contexts show the heterogeneity of the underlying records. **D)** Molecule classes are summarized across the packet. This figure supports Results Section 2.2 and Figure 2D; the authoritative row-level export is Table S3. The panels document reported-cargo provenance and sample heterogeneity; they do not measure preparation quality or EV specificity.

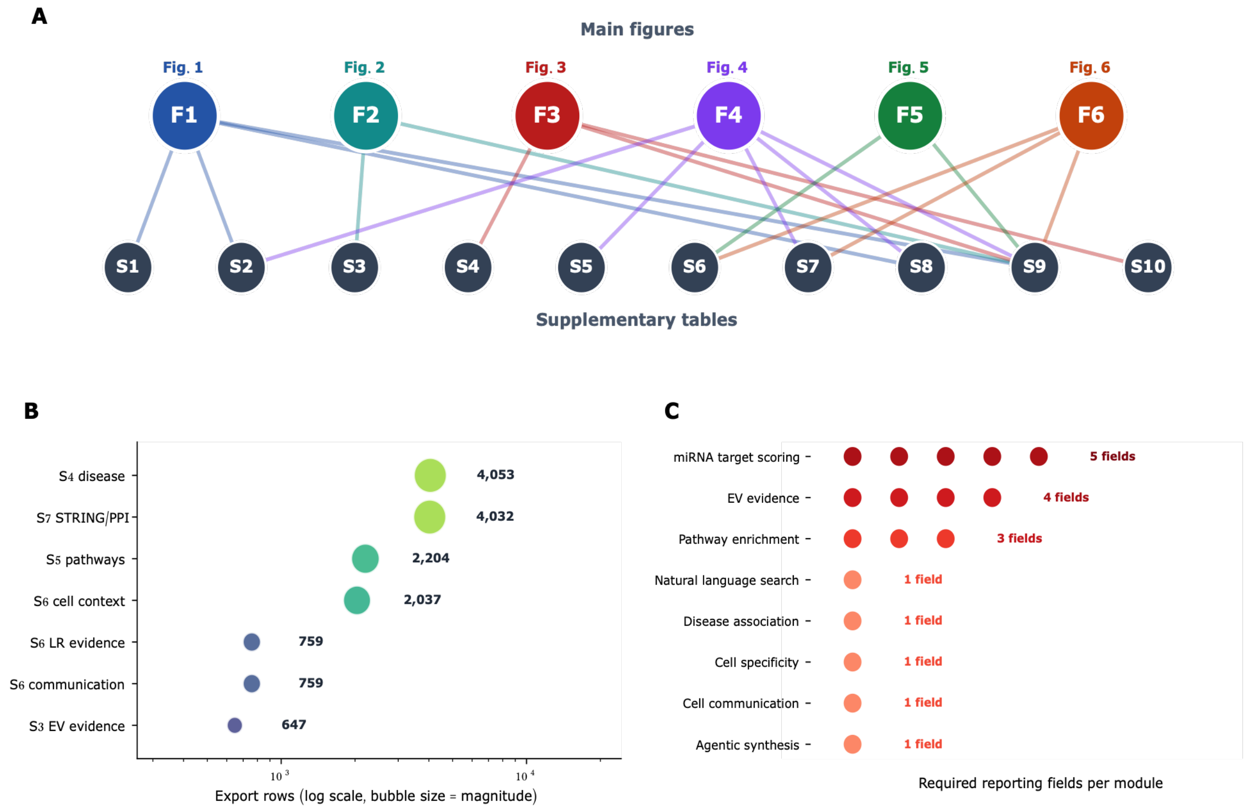

**Figure S4.** Links between main figures, supplementary exports, and evidence boundaries. **A)** Main Figures 1–6 are mapped to the exported evidence tables. **B)** Export row counts show the size of each evidence layer. **C)** Table S9 records the permitted interpretation, unsupported claims, and independent verification required before a stronger conclusion is made. This figure is the navigation map for the Results, Methods, and Supplementary Information.

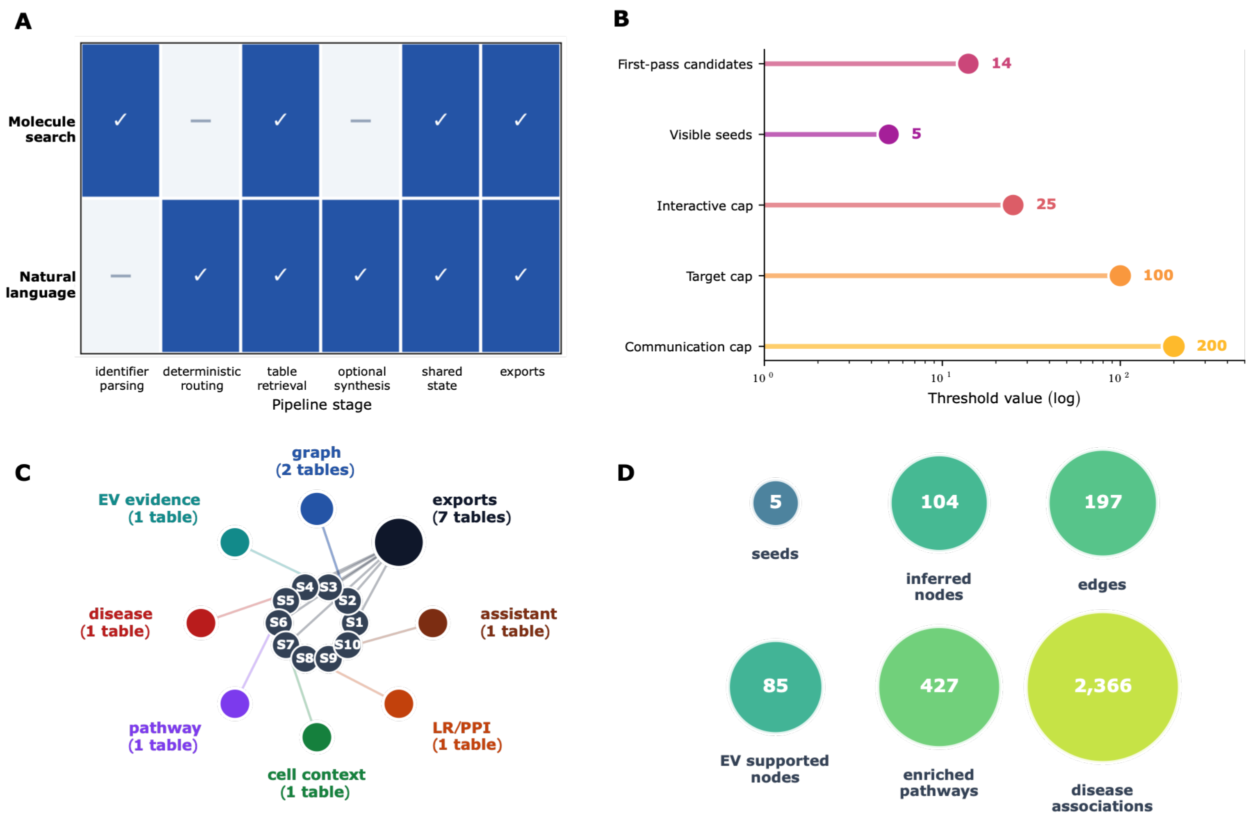

**Figure S5.** Explicit molecule search and natural-language routing converge on a shared evidence packet. **A)** Molecule and natural-language inputs enter distinct routing stages before table retrieval. **B)** The active processing guards are shown as computational limits. **C)** Supplementary-table depth is mapped across the analysis modules. **D)** The recorded case-study packet contains five visible starting molecules, 104 inferred molecules, 197 relationships, 85 nodes with reported EV support, 427 pathway summaries, and 2,366 disease summaries. This figure supports Results Section 2.1, Figure 1, and Methods: Query workflow and reviewable evidence packet and Natural-language navigation and grounding; the complete parameters are in Table S2. Processing caps are runtime limits, not biological thresholds.

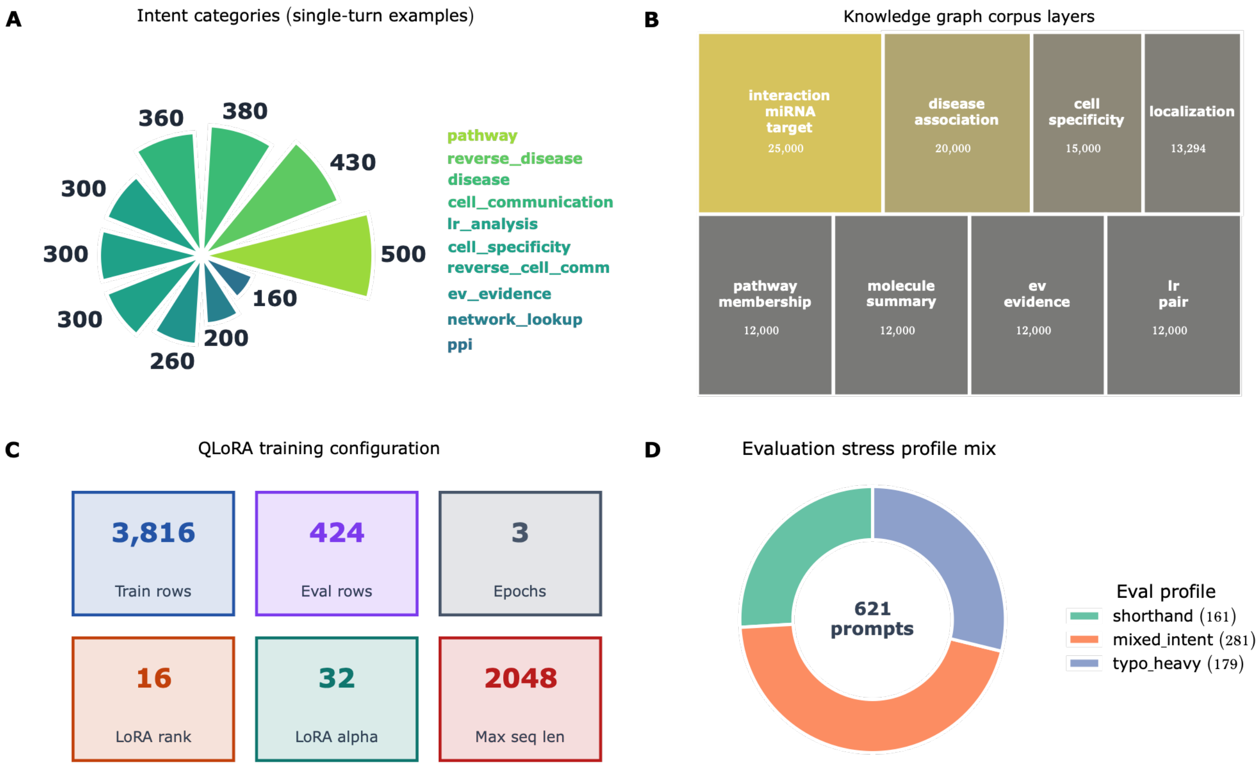

**Figure S6.** Current assistant training and grounding summary. **A)** Intent categories summarize the routing tasks represented in the released corpus. **B)** Knowledge-corpus layers show the table-backed evidence available to the assistant. **C)** QLoRA configuration records the principal fine-tuning parameters. **D)** The stress-prompt profile includes shorthand, mixed-intent, typo-heavy, and degraded inputs. This figure supports Figure 1G, Discussion Section 3.3, and Methods: Natural-language navigation and grounding; complete metadata are in Table S8. These records evaluate navigation over retrieved tables, not biological adjudication or EV function.

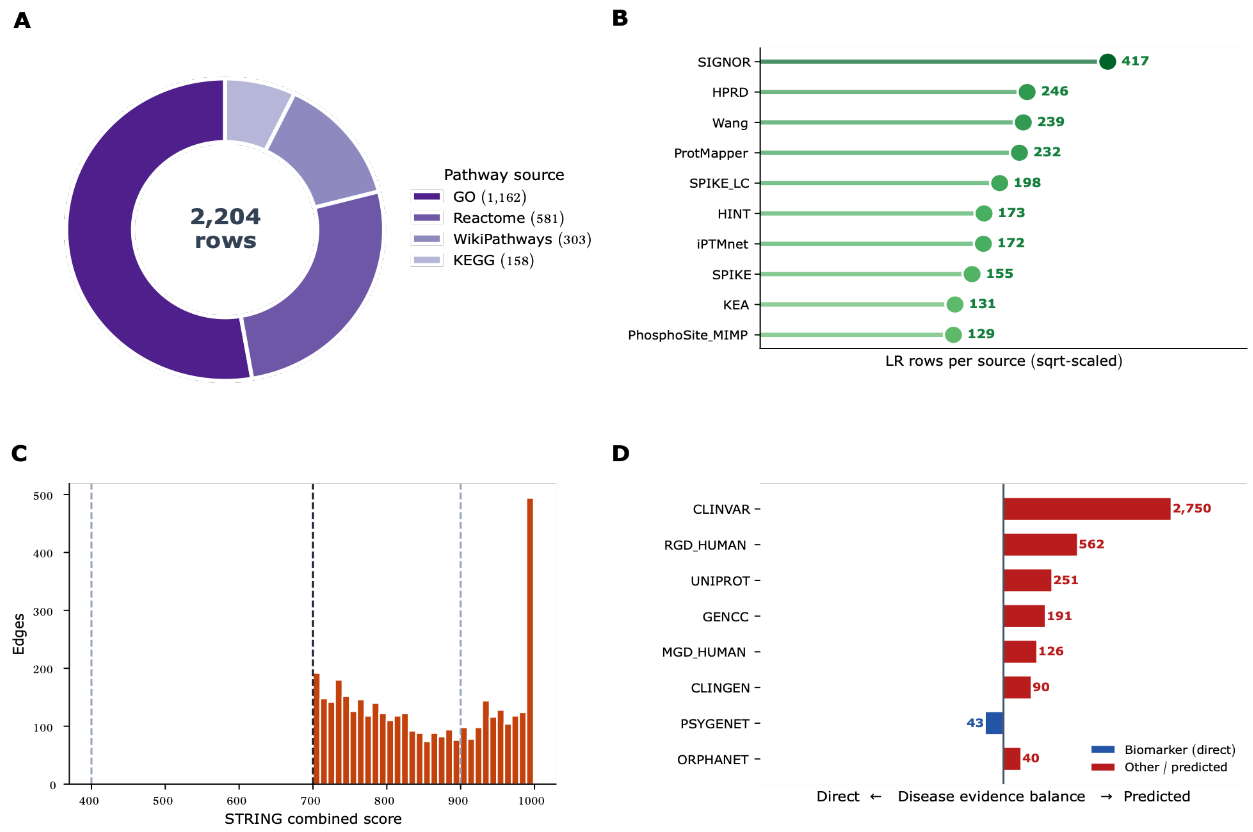

**Figure S7.** Source and threshold summaries for pathway, communication, PPI, and disease modules. **A)** Pathway-membership rows are summarized by source. **B)** Ligand–receptor rows are summarized by contributing source. **C)** The STRING combined-score distribution shows the high-confidence threshold used for review. **D)** Disease rows are separated by direct and predicted evidence classes. This figure supports Results Sections 2.2–2.6 and Methods: Disease and pathway annotation, Cell-context and communication prioritization, and miRNA-target and PPI bridge prioritization; row-level evidence is in Tables S4–S7. The panels make denominator, source, threshold, and literature-density dependence visible.

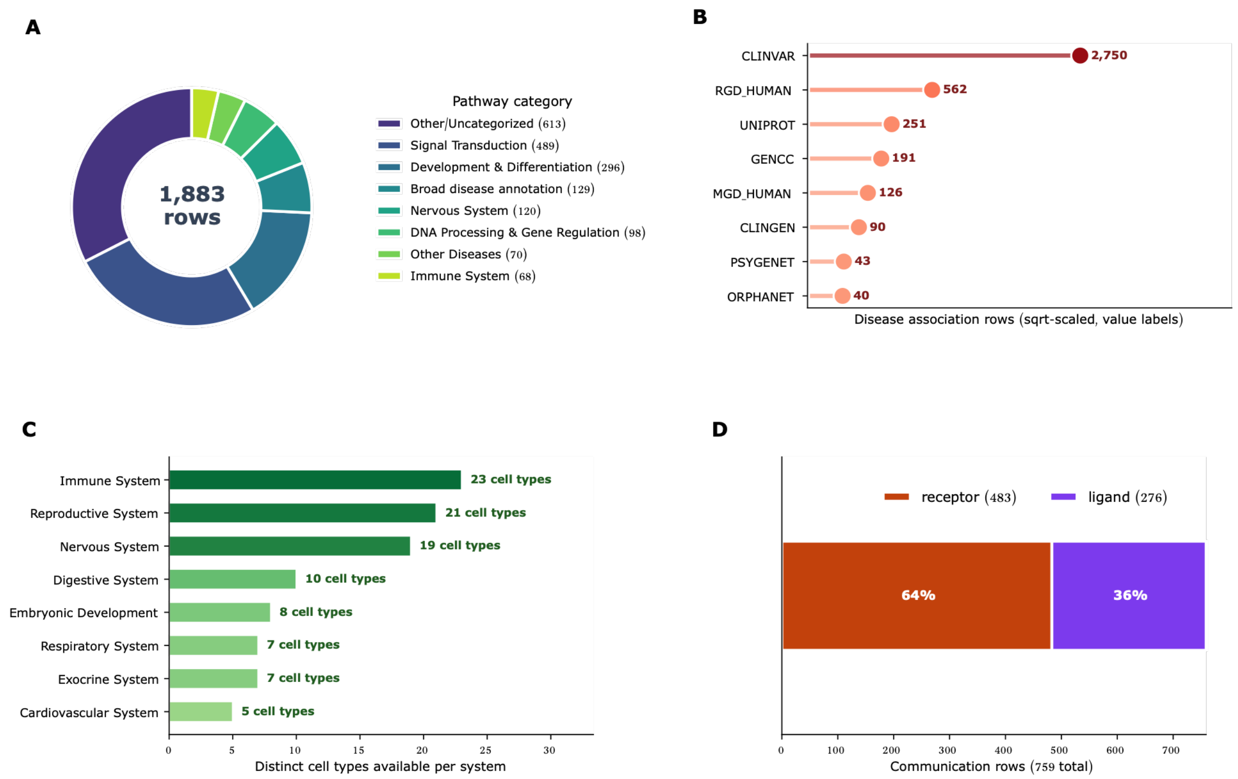

**Figure S8.** Cross-module audit summaries after EV-evidence review. **A)** Pathway memberships are grouped into broader categories. **B)** Disease-association rows are summarized by source. **C)** Available cell contexts are counted across biological systems. **D)** Communication candidates are summarized by ligand and receptor role. This figure supports Results Sections 2.3–2.5, Figures 3–5, and Methods: Disease and pathway annotation and Cell-context and communication prioritization; authoritative rows are in Tables S4–S6. The summaries support triage and expose source imbalance; they do not add EV-specific proof.

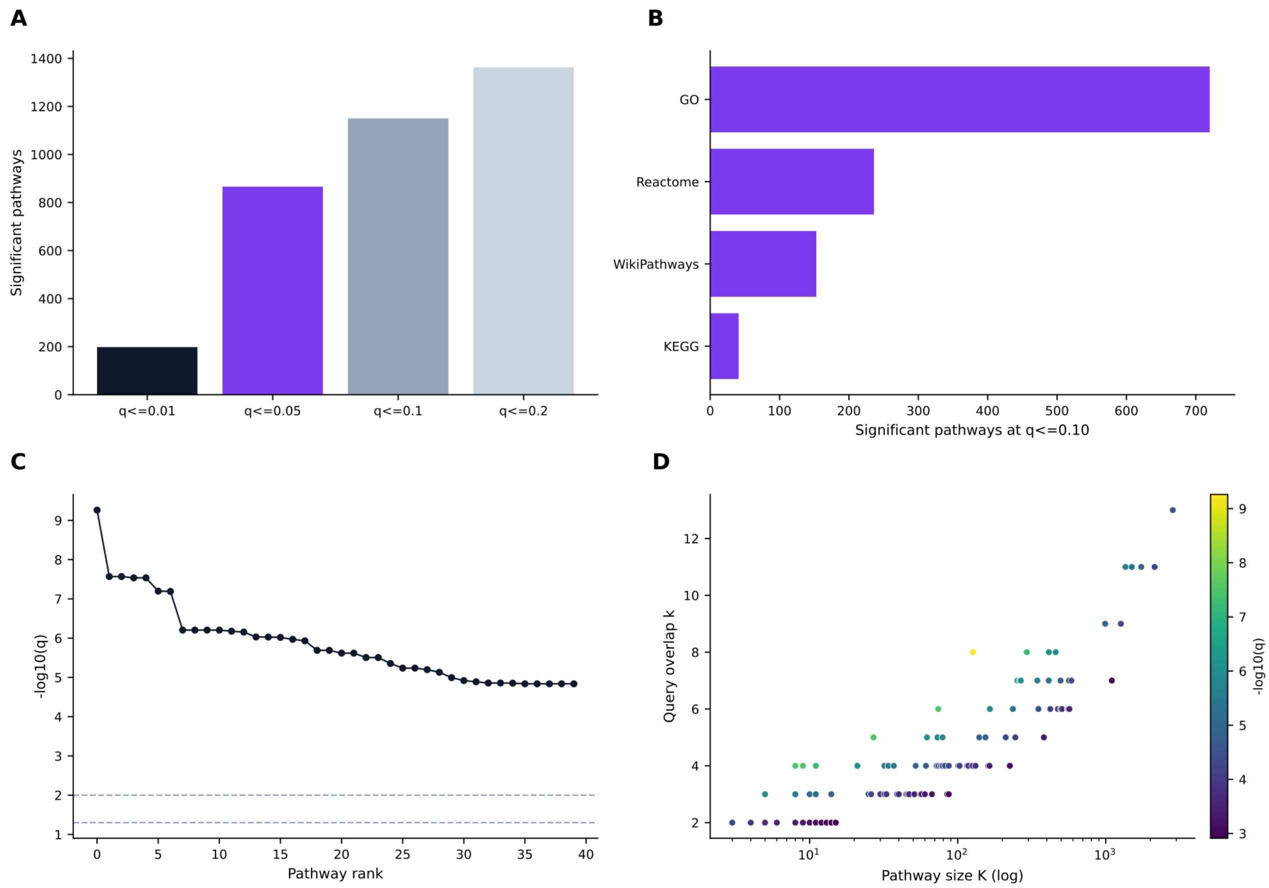

**Figure S9.** Pathway cutoff behavior from the exported pathway table. **A)** Significant-term counts are compared across q-value thresholds. **B)** Source-specific enrichment is shown at q ≤ 0.10. **C)** The highest-ranked pathway terms are ordered by adjusted significance. **D)** Query overlap is compared with pathway size. This figure supports Results Section 2.4, Figure 4, and Methods: Disease and pathway annotation; memberships and statistics are in Table S5. The panels document annotation-level enrichment behavior and do not measure pathway activity.

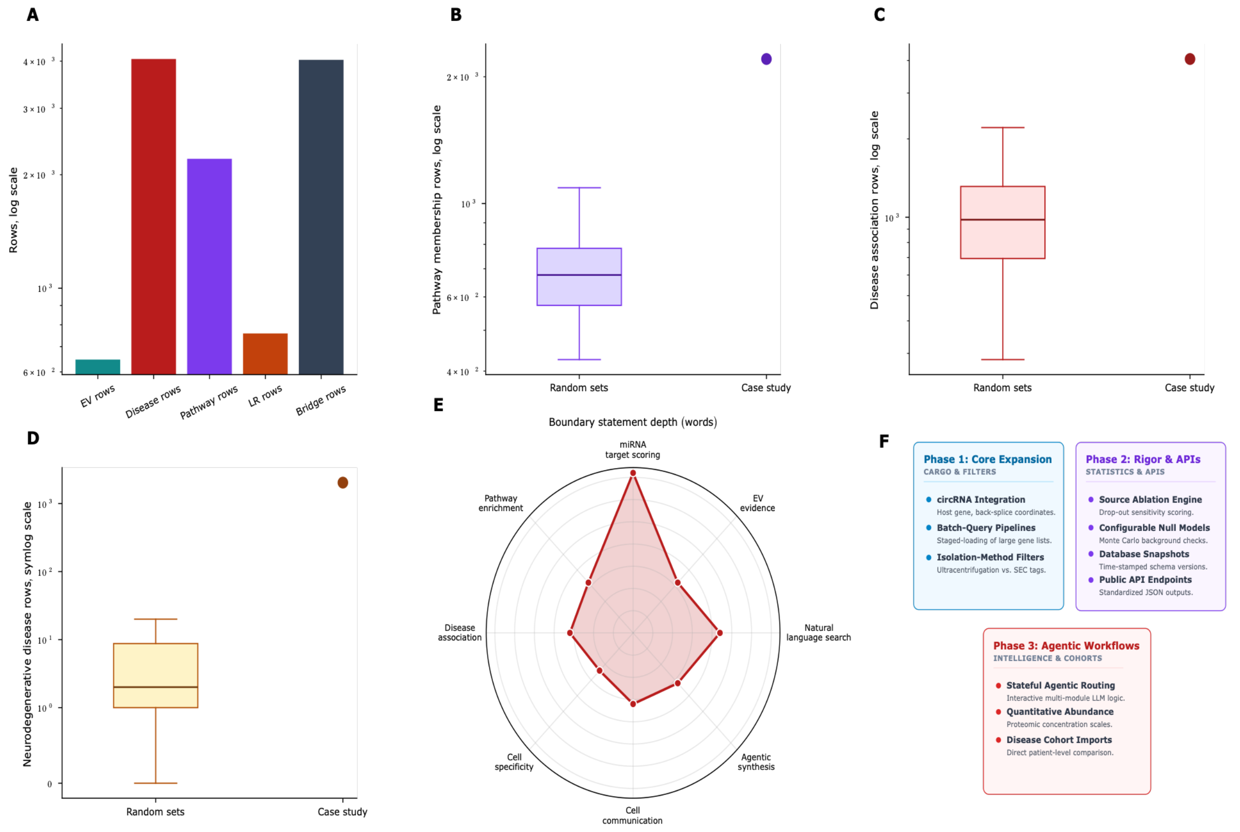

**Figure S10.** Sensitivity checks, evidence boundaries, and development roadmap. **A)** Evidence-layer row counts summarize the case-study exports. **B–D)** Random-set comparisons assess pathway, total disease, and neurodegenerative-disease annotation burden under the available database backgrounds. **E)** Table S9 summarizes the depth of module-specific boundary statements. **F)** The roadmap extends EV reporting and API functions toward an expert-reviewed claim–evidence training resource. Panels A–D support Results Section 2.6 and Methods: Sensitivity analysis and source-dependence checks; panels E–F support Discussion Sections 3.2–3.3 and Tables S9–S10. These panels evaluate database coherence and planned infrastructure, not EV-mediated mechanism.

#### 16 Supplementary Tables

The supplementary tables contain source rows, parameters, scores, and interpretation records used in the manuscript. Table S9 is the central evidence-boundary table and should be read alongside the module-specific exports.

**Table S1.** Source architecture and provenance for the EVd3x evidence-aware workflow. This table supports Figure 1, Figure S1, and Methods: Canonical table construction and source harmonization. It contains the source inventory, canonical analysis-table inventory, processing provenance, identifier harmonization, upstream source files, scripts, processing-stage counts, and where-used fields.

**Table S2.** Case-study query, visible starting molecules, inferred candidates, expansion parameters, and processing guards. This table supports Results Section 2.1, Figure 1, Figure S5, and Methods: Query workflow and reviewable evidence packet. It records the Alzheimer query, visible starting molecules, retained bridge miRNAs, first-pass candidates, expanded targets, network neighbors, thresholds, molecule caps, runtime limits, and the settings needed to reproduce the PSEN1-centered evidence packet.

**Table S3.** Reported EV-evidence export with source, sample, method, publication, and EV-TRACK-linked fields. This table is the authoritative export for Results Section 2.2, Figure 2D, Figures S2–S3, and Methods: EV evidence retrieval and reported-cargo provenance. Rows document reported cargo evidence and the provenance needed to judge relevance and define subsequent verification.

**Table S4.** Disease-context annotation export. This table supports Results Section 2.3, Figure 3, Figures S7D and S8B, and Methods: Disease and pathway annotation. It contains disease-association rows, supporting entities, sources, publication fields, and priority-score components used for context review and literature triage.

**Table S5.** Pathway annotation and enrichment export. This table supports Results Section 2.4, Figure 4, Figures S7A and S9, and Methods: Disease and pathway annotation. It contains pathway memberships, enrichment statistics, source labels, overlap genes, and cutoff-dependent records used to organize candidate biological branches for subsequent testing.

**Table S6.** Cell-context, communication, and ligand-receptor evidence export. This table supports Results Sections 2.5–2.6, Figures 5–6, Figures S7B and S8C–D, and Methods: Cell-context and communication prioritization. Rows retain plausible cell settings, communication pairs, molecular roles, expression context, score components, and source fields for review and independent verification.

**Table S7.** PPI and miRNA-to-mRNA bridge evidence export. This table supports Results Sections 2.2 and 2.6, Figures 2C and 6, Figure S7C, and Methods: miRNA-target and PPI bridge prioritization. It contains STRING/PPI rows and miRNA-target bridges used to prioritize, reject, or clarify molecular relationships before independent verification.

**Table S8.** Assistant corpus, deployment constraints, runtime candidates, and model metadata. This table supports Figure 1G, Discussion Sections 3.2–3.3, Figure S6, and Methods: Natural-language navigation and grounding. It contains training-corpus summaries, intent counts, evaluation profiles, adapter metadata, grounding policy, current limitations, and requirements for future EV evidence-training resources.

**Table S9.** Evidence boundaries and claim support by analysis layer. This table supports Table 1, Discussion Sections 3.1-3.2, Figure S4, and Figure S10E. It records permitted use, unsupported use, required verification, database-association limits, assistant-output limits, view-capping limits, API-reporting minimums, and main-text claim-to-evidence mapping.

**Table S10.** Future roadmap for EV reporting, reproducibility, database refreshes, API access, source snapshots, expert evidence labeling, AI benchmarking, and preservation of claim boundaries. This table supports Discussion Section 3.3 and Figure S10F. Planned items are not current manuscript claims.

#### References

[1] J. Lötvall, A. F. Hill, F. Hochberg, et al., “Minimal experimental requirements for definition of extracellular vesicles and their functions: A position statement from the International Society for Extracellular Vesicles,” Journal of Extracellular Vesicles, vol. 3, no. 1, p. 26913, 2014, doi: 10.3402/jev.v3.26913.

[2] C. Théry, K. W. Witwer, E. Aikawa, et al., “Minimal information for studies of extracellular vesicles 2018 (MISEV2018): A position statement of the International Society for Extracellular Vesicles and update of the MISEV2014 guidelines,” Journal of Extracellular Vesicles, vol. 7, no. 1, p. 1535750, 2018, doi: 10.1080/20013078.2018.1535750.

[3] J. A. Welsh, D. C. I. Goberdhan, L. O’Driscoll, et al., “Minimal information for studies of extracellular vesicles (MISEV2023): From basic to advanced approaches,” Journal of Extracellular Vesicles, vol. 13, no. 2, p. e12404, 2024, doi: 10.1002/jev2.12404.

[4] J. Van Deun, P. Mestdagh, P. Agostinis, et al., “EV-TRACK: Transparent reporting and centralizing knowledge in extracellular vesicle research,” Nature Methods, vol. 14, no. 3, pp. 228–232, 2017, doi: 10.1038/nmeth.4185.

[5] Q. Roux, J. Van Deun, S. Dedeyne, and A. Hendrix, “The EV-TRACK summary add-on: Integration of experimental information in databases to ensure comprehensive interpretation of biological knowledge on extracellular vesicles,” Journal of Extracellular Vesicles, vol. 9, no. 1, p. 1699367, 2020, doi: 10.1080/20013078.2019.1699367.

[6] B. Mateescu, E. J. K. Kowal, B. W. M. van Balkom, et al., “Obstacles and opportunities in the functional analysis of extracellular vesicle RNA–an ISEV position paper,” Journal of Extracellular Vesicles, vol. 6, no. 1, p. 1286095, 2017, doi: 10.1080/20013078.2017.1286095.

[7] R. Crescitelli, J. M. Falcón-Pérez, A. Hendrix, et al., “Reproducibility of extracellular vesicle research,” Journal of Extracellular Vesicles, vol. 14, no. 1, p. e70036, 2025, doi: 10.1002/jev2.70036.

[8] P. W. Harrison, M. R. Amode, O. Austine-Orimoloye, et al., “Ensembl 2024,” Nucleic Acids Research, vol. 52, no. D1, pp. D891–D899, 2024, doi: 10.1093/nar/gkad1049.

[9] The UniProt Consortium, “UniProt: The Universal Protein Knowledgebase in 2023,” Nucleic Acids Research, vol. 51, no. D1, pp. D523–D531, 2023, doi: 10.1093/nar/gkac1052.

[10] A. Kozomara and S. Griffiths-Jones, “miRBase: Annotating high confidence microRNAs using deep sequencing data,” Nucleic Acids Research, vol. 42, no. D1, pp. D68–D73, 2014, doi: 10.1093/nar/gkt1181.

[11] The RNAcentral Consortium, “RNAcentral 2021: Secondary structure integration, improved sequence search and new member databases,” Nucleic Acids Research, vol. 49, no. D1, pp. D212–D220, 2021, doi: 10.1093/nar/gkaa921.

[12] S. Mathivanan and R. J. Simpson, “ExoCarta: A compendium of exosomal proteins and RNA,” Proteomics, vol. 9, no. 21, pp. 4997–5000, 2009, doi: 10.1002/pmic.200900351.

[13] S. V. Chitti, S. Gummadi, T. Kang, et al., “Vesiclepedia 2024: An extracellular vesicles and extracellular particles repository,” Nucleic Acids Research, vol. 52, no. D1, pp. D1694–D1698, 2024, doi: 10.1093/nar/gkad1007.

[14] Z. Wei, N. Zhou, M. Jing, et al., “SVAtlas: A comprehensive single extracellular vesicle omics resource,” Nucleic Acids Research, vol. 54, no. D1, pp. D1807–D1816, 2026, doi: 10.1093/nar/gkaf1189.

[15] H.-Y. Huang, Y.-C.-D. Lin, S. Cui, et al., “miRTarBase update 2022: An informative resource for experimentally validated miRNA-target interactions,” Nucleic Acids Research, vol. 50, no. D1, pp. D222–D230, 2022, doi: 10.1093/nar/gkab1079.

[16] D. Karagkouni, M. D. Paraskevopoulou, S. Chatzopoulos, et al., “DIANA-TarBase v8: A decade-long collection of experimentally supported miRNA-gene interactions,” Nucleic Acids Research, vol. 46, no. D1, pp. D239–D245, 2018, doi: 10.1093/nar/gkx1141.

[17] V. Agarwal, G. W. Bell, J.-W. Nam, and D. P. Bartel, “Predicting effective microRNA target sites in mammalian mRNAs,” eLife, vol. 4, p. e05005, 2015, doi: 10.7554/eLife.05005.

[18] M. Gillespie, B. Jassal, R. Stephan, et al., “The Reactome pathway knowledgebase 2022,” Nucleic Acids Research, vol. 50, no. D1, pp. D687–D692, 2022, doi: 10.1093/nar/gkab1028.

[19] M. Kanehisa, M. Furumichi, Y. Sato, M. Kawashima, and M. Ishiguro-Watanabe, “KEGG for taxonomy-based analysis of pathways and genomes,” Nucleic Acids Research, vol. 51, no. D1, pp. D587–D592, 2023, doi: 10.1093/nar/gkac963.

[20] The Gene Ontology Consortium, “The Gene Ontology knowledgebase in 2023,” Genetics, vol. 224, no. 1, p. iyad031, 2023, doi: 10.1093/genetics/iyad031.

[21] M. Martens, A. Ammar, A. Riutta, et al., “WikiPathways: Connecting communities,” Nucleic Acids Research, vol. 49, no. D1, pp. D613–D621, 2021, doi: 10.1093/nar/gkaa1024.

[22] J. Piñero, J. M. Ramírez-Anguita, J. Saüch-Pitarch, et al., “The DisGeNET knowledge platform for disease genomics: 2019 update,” Nucleic Acids Research, vol. 48, no. D1, pp. D845–D855, 2020, doi: 10.1093/nar/gkz1021.

[23] P. J. Thul and C. Lindskog, “The Human Protein Atlas: A spatial map of the human proteome,” Protein Science, vol. 27, no. 1, pp. 233–244, 2018, doi: 10.1002/pro.3307.

[24] T. Cui, Y. Dou, P. Tan, et al., “RNALocate v2.0: An updated resource for RNA subcellular localization with increased coverage and annotation,” Nucleic Acids Research, vol. 50, no. D1, pp. D333–D339, 2022, doi: 10.1093/nar/gkab825.

[25] A. Keller, L. Gröger, T. Tschernig, et al., “miRNATissueAtlas2: An update to the human miRNA tissue atlas,” Nucleic Acids Research, vol. 50, no. D1, pp. D211–D221, 2022, doi: 10.1093/nar/gkab808.

[26] B. Panwar, G. S. Omenn, and Y. Guan, “miRmine: A database of human miRNA expression profiles,” Bioinformatics, vol. 33, no. 10, pp. 1554–1560, 2017, doi: 10.1093/bioinformatics/btx019.

[27] X. Zhang, Y. Lan, J. Xu, et al., “CellMarker: A manually curated resource of cell markers in human and mouse,” Nucleic Acids Research, vol. 47, no. D1, pp. D721–D728, 2019, doi: 10.1093/nar/gky900.

[28] O. Franzén, L.-M. Gan, and J. L. M. Björkegren, “PanglaoDB: A web server for exploration of mouse and human single-cell RNA sequencing data,” Database, vol. 2019, p. baz046, 2019, doi: 10.1093/database/baz046.

[29] R. Vento-Tormo, M. Efremova, R. A. Botting, et al., “Single-cell reconstruction of the early maternal-fetal interface in humans,” Nature, vol. 563, no. 7731, pp. 347–353, 2018, doi: 10.1038/s41586-018-0698-6.

[30] M. Efremova, M. Vento-Tormo, S. A. Teichmann, and R. Vento-Tormo, “CellPhoneDB: Inferring cell-cell communication from combined expression of multi-subunit ligand-receptor

complexes," Nature Protocols, vol. 15, no. 4, pp. 1484–1506, 2020, doi: 10.1038/s41596-020-0292-x.

[31] D. Türei, A. Valdeolivas, L. Gul, et al., "Integrated intra- and intercellular signaling knowledge for multicellular omics analysis," Molecular Systems Biology, vol. 17, no. 3, p. e9923, 2021, doi: 10.15252/msb.20209923.

[32] X. Shao, J. Liao, C. Li, X. Lu, J. Cheng, and X. Fan, "CellTalkDB: A manually curated database of ligand-receptor interactions in humans and mice," Briefings in Bioinformatics, vol. 22, no. 4, p. bbaa269, 2021, doi: 10.1093/bib/bbaa269.

[33] Y. Zhang, T. Liu, J. Wang, et al., "Cellinker: A platform of ligand-receptor interactions for intercellular communication analysis," Bioinformatics, vol. 37, no. 14, pp. 2025–2032, 2021, doi: 10.1093/bioinformatics/btab036.

[34] D. Szklarczyk, R. Kirsch, M. Koutrouli, et al., "The STRING database in 2023: Protein-protein association networks and functional enrichment analyses for any sequenced genome of interest," Nucleic Acids Research, vol. 51, no. D1, pp. D638–D646, 2023, doi: 10.1093/nar/gkac1000.

[35] Y. Benjamini and Y. Hochberg, "Controlling the false discovery rate: A practical and powerful approach to multiple testing," Journal of the Royal Statistical Society: Series B (Methodological), vol. 57, no. 1, pp. 289–300, 1995, doi: 10.1111/j.2517-6161.1995.tb02031.x.

[36] E. J. Hu et al., "LoRA: Low-rank adaptation of large language models," arXiv preprint arXiv:2106.09685, 2021, doi: 10.48550/arXiv.2106.09685.

[37] T. Dettmers, A. Pagnoni, A. Holtzman, and L. Zettlemoyer, "QLoRA: Efficient finetuning of quantized LLMs," Advances in Neural Information Processing Systems, vol. 36, pp. 10088–10115, 2023, doi: 10.48550/arXiv.2305.14314.

[38] J. Bai et al., "Qwen technical report," arXiv preprint arXiv:2309.16609, 2023, doi: 10.48550/arXiv.2309.16609.

[39] Qwen Team, "Qwen2.5 technical report," arXiv preprint arXiv:2412.15115, 2024, doi: 10.48550/arXiv.2412.15115.

[40] S. Memczak, M. Jens, A. Elefsinioti, et al., "Circular RNAs are a large class of animal RNAs with regulatory potency," Nature, vol. 495, no. 7441, pp. 333–338, 2013, doi: 10.1038/nature11928.

[41] L.-L. Chen, "The biogenesis and emerging roles of circular RNAs," Nature Reviews Molecular Cell Biology, vol. 17, no. 4, pp. 205–211, 2016, doi: 10.1038/nrm.2015.32.

[42] Y. Li, Q. Zheng, C. Bao, et al., "Circular RNA is enriched and stable in exosomes: A promising biomarker for cancer diagnosis," Cell Research, vol. 25, no. 8, pp. 981–984, 2015, doi: 10.1038/cr.2015.82.
